# SAMbinder: A web server for predicting SAM binding residues of a protein from its amino acid sequence

**DOI:** 10.1101/625806

**Authors:** Piyush Agrawal, Gaurav Mishra, Gajendra P. S. Raghava

## Abstract

**Motivation:** S-adenosyl-L-methionine (SAM) is one of the important cofactor present in the biological system and play a key role in many diseases. There is a need to develop a method for predicting SAM binding sites in a protein for designing drugs against SAM associated disease. Best of our knowledge, there is no method that can predict the binding site of SAM in a given protein sequence.

**Result:** This manuscript describes a method SAMbinder, developed for predicting SAM binding sites in a protein from its primary sequence. All models were trained, tested and evaluated on 145 SAM binding protein chains where no two chains have more than 40% sequence similarity. Firstly, models were developed using different machine learning techniques on a balanced dataset contain 2188 SAM interacting and an equal number of non-interacting residues. Our Random Forest based model developed using binary profile feature got maximum MCC 0.42 with AUROC 0.79 on the validation dataset. The performance of our models improved significantly from MCC 0.42 to 0.61, when evolutionary information in the form of PSSM profile is used as a feature. We also developed models on realistic dataset contains 2188 SAM interacting and 40029 non-interacting residues and got maximum MCC 0.61 with AUROC of 0.89. In order to evaluate the performance of our models, we used internal as well as external cross-validation technique.

**Availability and implementation:** https://webs.iiitd.edu.in/raghava/sambinder/.

## Introduction

Structural and functional annotation of a protein is one among the major challenges in the era of genomics. With the rapid advancement in sequencing technologies and concerted genome projects, there is an increasing gap between the sequenced protein and functionally annotated proteins which are very few in number (Casari *et al*., 1995; Yu *et al*., 2014). Therefore, there is an urgent need for the development of automated computational methods which can identify the residues playing an important role in protein functions. Protein-ligand interaction has been identified as one of the important functions which play a vital function in all biological processes (Agrawal, Singh, *et al*., 2019). In the past, considerable efforts have been to develop tools which can identify the ligand binding residues in a protein (Sousa *et al*., 2006). In the early stage, generalized methods have been developed which predicts the binding site or pockets in the proteins regardless of their ligand (Hendlich *et al*., 1997; Dundas *et al*., 2006; Laskowski, 1995; Levitt and Banaszak, 1992; Le Guilloux *et al*., 2009). Later on, it was realized that all ligands are not the same and there is a wide variation in shape and size of binding pockets. Therefore, researchers started developing ligand specific methods (Chauhan, Mishra, and G. P. S. Raghava, 2009; Hu *et al*., 2018; Yu, Hu, Huang, *et al*., 2013; Chen *et al*., 2012; Chauhan *et al*., 2010; Hu *et al*., 2016), and it was observed that these ligand specific methods performed better than generalized methods (Yu, Hu, Yang, *et al*., 2013a; Chen *et al*., 2012; Hu *et al*., 2016).

All living organism consists of small molecular weight ligands or cofactors which carries out an important function in some metabolic and regulatory pathways. S-adenosyl-L-methionine (SAM) is one such important cofactor, which was first discovered in the year 1952. After ATP, SAM is the second most versatile and widely used small molecule (Cantoni, 1975). It is a natural substance present in the cells of the body and is a direct metabolite of L-methionine which is an essential amino acid. SAM is a conjugate molecule of two ubiquitous biological compounds; (i) adenosine moiety of ATP and (ii) amino acid methionine (CATONI, 1953; Waddell *et al*., 2000). One of the most important functions of the SAM is the transfer or donation of different chemical groups such as methyl (Thomas *et al*., 2004; Wuosmaa and Hager, 1990), aminopropyl (Lin, 2011), ribosyl (Kozbial and Mushegian, 2005), 5’deocxyadenosyl and methylene group (Gana *et al*., 2013; Kozbial and Mushegian, 2005) for carrying out covalent modification of a variety of substrates. SAM is also used as a precursor molecule in the biosynthesis of nicotinamide phytosiderophores, plant hormone ethylene, spermine, and spermidine; carry out chemical reactions such as hydroxylation, fluorination which takes place in bacteria (Cadicamo *et al*., 2004). It has become the choice of various clinical studies and possess therapeutic value for treating diseases like osteoarthritis (Najm *et al*., 2004), cancer (Chaib *et al*., 2011; Wagner *et al*., 2010), epilepsy (Item *et al*., 2004), Alzheimer’s (Borroni *et al*., 2004), dementia and depression (Bottiglieri *et al*., 1990; Rosenbaum *et al*., 1990), Parkinson (Zhu, 2004), and other psychiatric and neurological disorders (Bottiglieri, 1997). In the previous studies, it has been shown that mutation in the binding site of SAM has changed the protein function. For example, Aktas et. al. showed that Alanine substitution in the predicted SAM binding residues reduced the SAM binding affinity and enzyme activity dramatically (Aktas *et al*., 2011). Thus, there is a need to develop a method that can predict SAM binding sites in a protein as it is an important ligand.

## Material and Methods

### Dataset Creation

In the first step, we extracted 244 SAM binding proteins PDB IDs from ccPDB 2.0 (Agrawal, Patiyal, *et al*., 2019). In total, we obtained 457 SAM binding protein chains. In the next step, we filtered all the sequences with 40% sequence similarity using CD-HIT software (Huang *et al*., 2010) and resolution better than 3 Å and obtained total of 145 protein chains. In the next step, we run Ligand Protein Contact (LPC) software (Sobolev *et al*., 1999) on the chains to extract the contact information of SAM with residues present in the protein chains with cutoff criteria of 4 Å, which means if the distance of contact of SAM with a residue is less than or equal to 4 Å, the residue is called interacting otherwise the residue is called as non-interacting. This is well-established criteria adopted in many previous studies (Chauhan *et al*., 2010; Mishra and Raghava, 2010).

### Additional dataset creation

An additional dataset was also created consisting of the SAM binding chains released after March 2018 and up to February 28, 2019. The dataset was generated following the same procedures as mentioned above and comprised of total 10 chains.

### Internal and External Validation

Dataset was randomly divided into two parts; (i) training dataset, which comprises 80% of the protein chains and (ii) validation dataset, which consists of remaining 20% of the protein chains. Datasets were created at the protein level and not at pattern or residue level since in previous studies it has been shown that dataset generated at pattern level is biased and show higher performance (Yu *et al*., 2014). The balanced dataset contains 1798 SAM interacting and non-interacting residue in training dataset and 390 SAM interacting and non-interacting residue in the validation dataset. The realistic dataset consists of 1798 SAM interacting residues and 33314 SAM non-interacting residues in training dataset whereas validation dataset consists of 390 SAM interacting residues and 6715 SAM non-interacting residues.

### Five-fold cross-validation

The five-fold cross technique was performed for evaluating the performance of different prediction models. This kind of performance evaluation has been used in many previous studies (Nagpal *et al*., 2018; Kumar *et al*., 2018).

### Window or Pattern size

We created overlapping patterns of each sequence of different window size ranging from 5-23 amino acid length. If the pattern central residue is SAM interacting, it is assigned as a positive pattern; otherwise, it was assigned as a negative pattern. In order to generate the pattern for terminus residues, we added (L-1)/2 dummy residue ‘X’ at both the termini of the protein chain (where L is pattern length).

### Binary Profile

We generated binary profile of each patterns by assigning binary values to the amino acids in fixed length pattern. A vector of dimension 21 represented each amino acid present in the pattern hence leading to final vector of N x 21, where N is length of the pattern. For example, residue ‘A’ was represented by 1,0,0,0,0,0,0,0,0,0,0,0,0,0,0,0,0,0,0,0,0; which contains 20 amino acids and one dummy amino acid ‘X’. X was represented by 0,0,0,0,0,0,0,0,0,0,0,0,0,0,0,0,0,0,0,0,0 (Agrawal, Kumar, *et al*., 2019; Agrawal and Raghava, 2018).

### Position Specific Scoring Matrix (PSSM)

PSSM profiles containing evolutionary information has been shown as an important feature in many previous studies for predicting protein-ligand interaction and other bioinformatics problems (Yu, Hu, Huang, *et al*., 2013; Chen *et al*., 2012). PSSM profiles of a sequence were generated using Position-Specific Iterative Basic Local Alignment Search Tool (PSI-BLAST) and searching against the Swiss Prot database. Three iterations were performed with E-value cut-off of 0.001 against each sequence. The original PSSM profiles obtained were further normalized to get value in between 0 and 1, followed by calculation of position specific score for each residue. The final matrix obtained consists of 21× N elements (20 amino acids residue and one dummy residue ‘X’). Here N, is the length of the pattern.

### Machine Learning Techniques

We implemented the python based machine learning package SCIKIT-learn (Pedregosa FABIANPEDREGOSA *et al*., 2011) for developing prediction models. We implemented Support Vector Classifier (SVC), Random Forest classifier (RFC), ExtraTree classifier (ETC), K-Nearest Neighbor (KNN), Multilayer Perceptron (MLP) and Ridge classifier for developing prediction models. Before, developing prediction models, we optimized different parameters on our internal dataset using Grid Search parameter present in the package.

### Evaluation Parameters

Performance of developed prediction models was evaluated in terms of Sensitivity (Sen), Specificity (Spc), Accuracy (Acc), Matthew’s Correlation Coefficient (MCC) and Area Under Receiver Operating Characteristics (AUROC). ‘pROC package’ implemented in R was used for computing AUROC (Title Display and Analyze ROC Curves, 2019). The formula for calculating is explained in the equation 1-4.

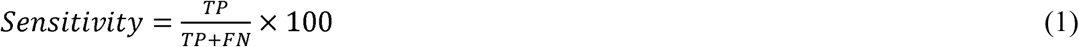

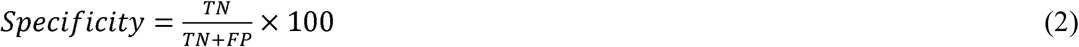

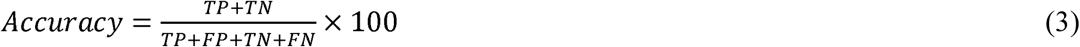

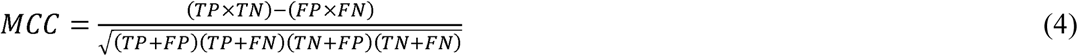

Where TP represents correctly predicted positive value, TN represents the correctly predicted negative value, FP represents the actual negative value which has been wrongly predicted as positive, and FN represents the positive value which has been wrongly predicted as negative.

## Results

### Composition Analysis

We have analyzed the amino acid composition of SAM interacting and non-interacting residues in SAM binding proteins. As shown in Figure 1, composition of residues C, D, F, G, H, M, N, S, W, and Y are higher in SAM interacting sites whereas composition of residues like A, E, I, K, L, P, Q, R, and V are higher in SAM non-interacting sites.

**Figure 1.**
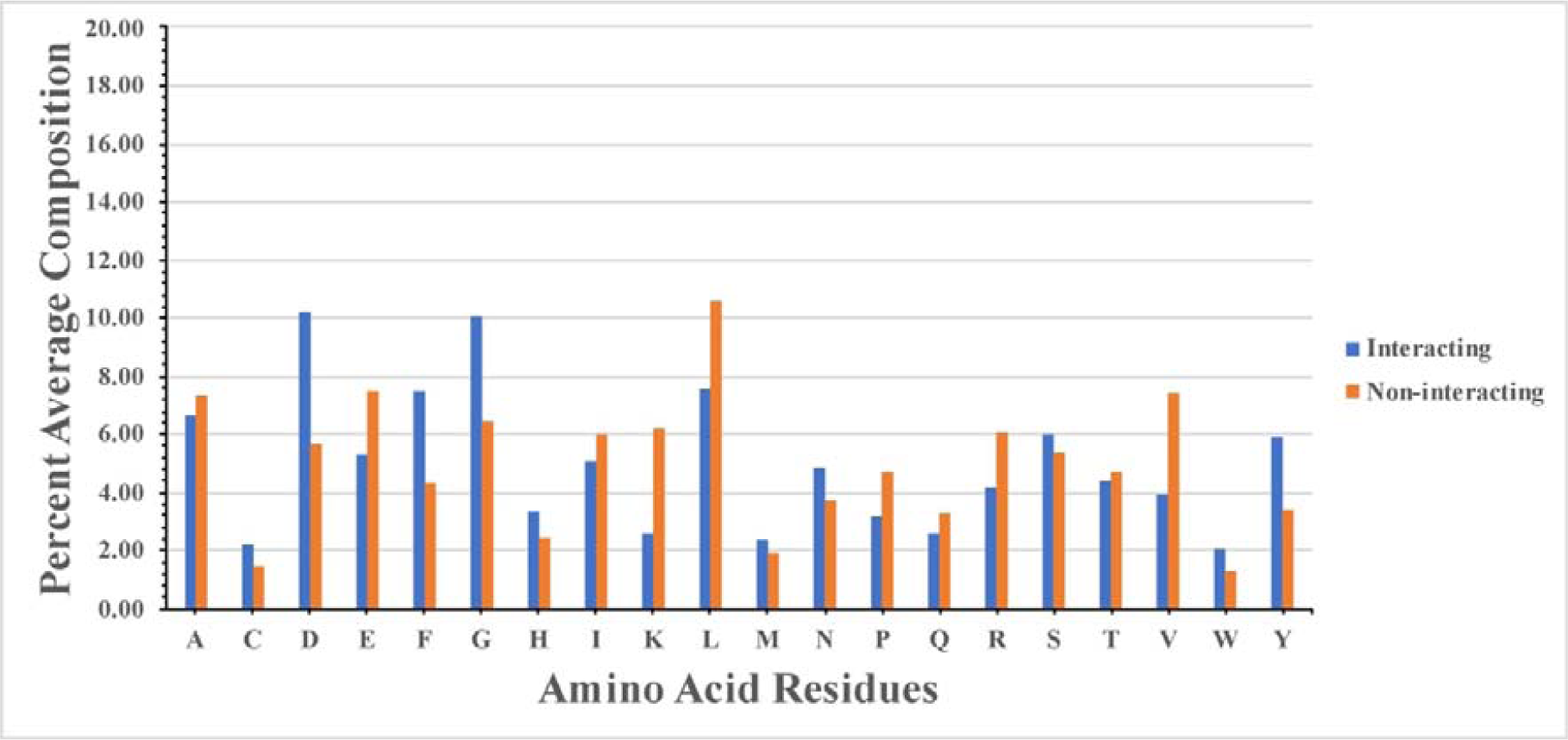
Percentage composition of SAM interacting and non-interacting residues.

### Propensity Analysis

We also analyzed the propensities of amino acid residues in SAM interacting and non-interacting sites. We observed that propensities of residues like C, D, F, G, H, M, N, S, W, and Y was higher in SAM interacting sites whereas propensities of residues like A, E, I, K, L, P, Q, R, T, and V was higher in SAM non-interacting sites (Supplementary Figure S1).

### Physiochemical Properties Analysis

We also analyzed the various physiochemical properties in SAM binding proteins. We found that SAM interacting sites are rich in acidic, small, polar and aromatic amino acids whereas SAM non-interacting sites are more predominant in charge, basic, and aliphatic amino acids. (Supplementary Figure S2).

### Machine learning model performance using binary patterns

Various machine learning models were developed using binary patterns for window size 5-23 on the balanced dataset. We compiled the best result obtained for each window size in Table 1 and plot the AUROC obtained for both training and validation dataset as shown in **Figure 2(a) and Figure 2(b)** respectively. In our analysis, we found that prediction model developed using random forest on the window size 21 performed best among all the prediction models. The model achieved accuracy of 70.79%, 0.42 MCC and 0.78 AUROC on the training dataset and accuracy of 70.85%, 0.42 MCC and 0.79 AUROC on the validation dataset. Detail result obtained for each window size by different machine learning techniques is provided in the **Supplementary Table S1-S10**.

**Table 1.** The performance of best machine learning model developed using amino acid sequence (binary pattern) for individual window size on balanced dataset.

|  | Training Dataset |  |  |  |  | Validation Dataset |  |  |  |  |
| --- | --- | --- | --- | --- | --- | --- | --- | --- | --- | --- |
|  | Sen | Spc | Acc | MCC | AUROC | Sen | Spc | Acc | MCC | AUROC |
| Pat5 (SVC) | 64.98 | 64.31 | 64.65 | 0.29 | 0.70 | 65.54 | 68.65 | 67.10 | 0.34 | 0.73 |
| Pat7 (RF) | 69.87 | 64.31 | 67.09 | 0.34 | 0.74 | 69.43 | 66.58 | 68.01 | 0.36 | 0.74 |
| Pat9 (RF) | 72.05 | 65.66 | 68.86 | 0.38 | 0.76 | 69.95 | 68.65 | 69.30 | 0.39 | 0.76 |
| Pat11 (RF) | 69.19 | 70.26 | 69.73 | 0.39 | 0.77 | 65.03 | 71.76 | 68.39 | 0.37 | 0.76 |
| Pat13 (RF) | 73.06 | 66.05 | 69.56 | 0.39 | 0.77 | 70.98 | 65.03 | 68.01 | 0.36 | 0.77 |
| Pat15 (RF) | 69.58 | 70.71 | 70.15 | 0.40 | 0.78 | 66.06 | 72.02 | 69.04 | 0.38 | 0.78 |
| Pat17 (RF) | 70.37 | 71.10 | 70.74 | 0.41 | 0.78 | 67.36 | 71.50 | 69.43 | 0.39 | 0.78 |
| Pat19 (RF) | 70.54 | 71.32 | 70.93 | 0.42 | 0.78 | 67.36 | 73.83 | 70.60 | 0.41 | 0.79 |
| Pat21 (RF) | 70.37 | 71.21 | 70.79 | 0.42 | 0.78 | 67.62 | 74.09 | 70.85 | 0.42 | 0.79 |
| Pat23 (RF) | 70.76 | 70.99 | 70.88 | 0.42 | 0.78 | 68.39 | 71.76 | 70.08 | 0.40 | 0.79 |

\* **Sen:** Sensitivity, **Spc:** Specificity, **Acc:** Accuracy, **MCC:** Matthews Correlation Coefficient, **AUROC:** Area Under the Receiver Operating Characteristic curve, **SVC:** Support Vector Classifier, **RF:** Random Forest.

**Figure 2.**
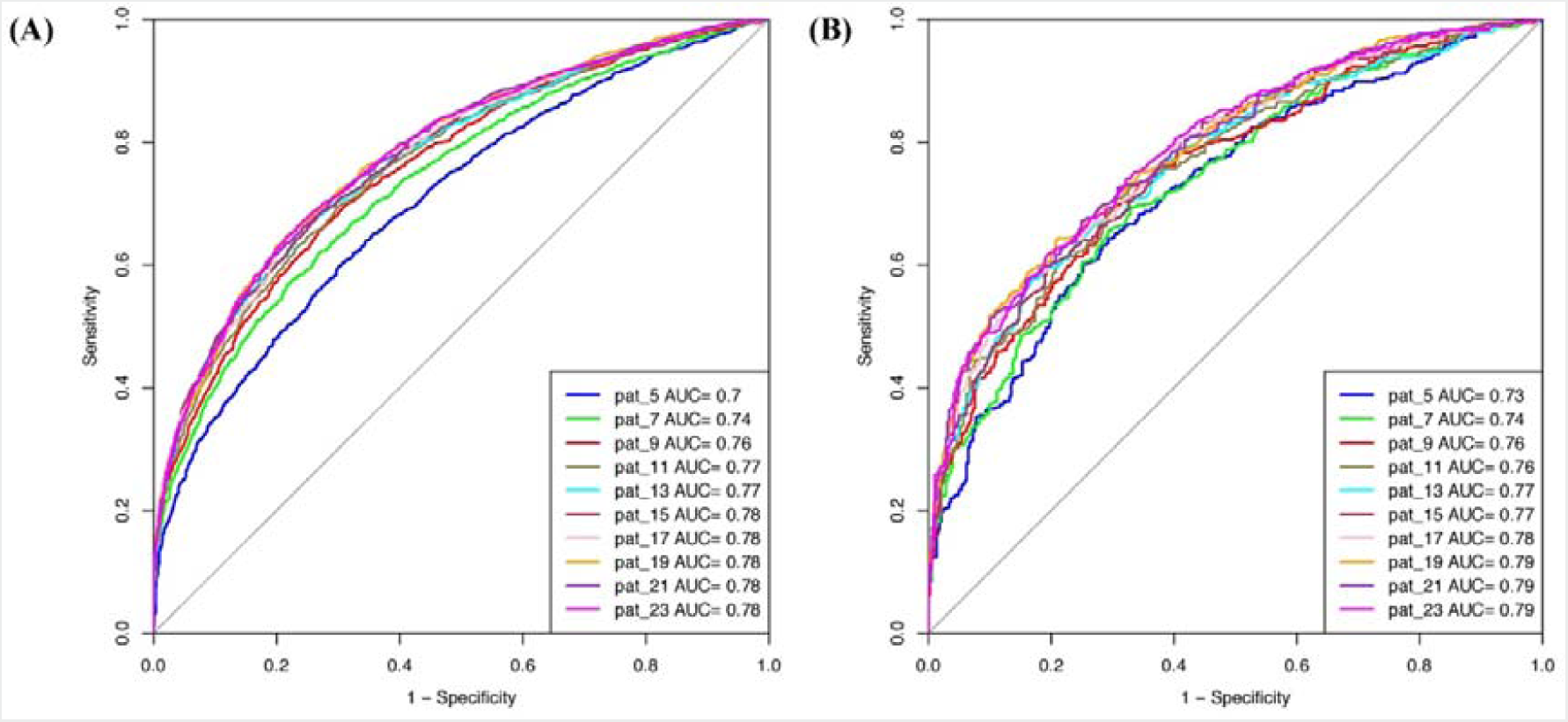
AUROC plots obtained for various window length developed using binary profile on balanced dataset for (a) training dataset and (b) validation dataset.

### Machine learning model performance using evolutionary information (PSSM profile)

Prediction models were developed using PSSM profiles for all the considered window size on the balanced dataset. Best result obtained for each window size is compiled in Table 2 and AUROC was plotted for the training **(Figure 3(a))** and validation dataset **(Figure 3(b))**. We observed that, in case of PSSM profiles, the performance of the prediction models were increased. ExtraTree Classifier model developed on the window size 17 performed best among all the developed models. It achieved the highest accuracy of 80.39%, MCC of 0.61 and AUROC of 0.88 on training dataset whereas on validation dataset it achieved accuracy of 77.07%, MCC of 0.54, and AUROC of 0.86. Result obtained by different classifiers on each window size has been provided in the **Supplementary Table S11-S20.**

**Table 2.** The performance of best machine learning model developed using PSSM profile for individual window size on balanced dataset.

|  | Training Dataset |  |  |  |  | Validation Dataset |  |  |  |  |
| --- | --- | --- | --- | --- | --- | --- | --- | --- | --- | --- |
|  | Sen | Spc | Acc | MCC | AUROC | Sen | Spc | Acc | MCC | AUROC |
| Pat5 (RF) | 78.40 | 77.50 | 77.95 | 0.56 | 0.86 | 74.35 | 76.94 | 75.65 | 0.51 | 0.83 |
| Pat7 (ETree) | 79.85 | 79.12 | 79.49 | 0.59 | 0.87 | 73.32 | 79.53 | 76.42 | 0.53 | 0.85 |
| Pat9 (ETree) | 81.48 | 76.49 | 78.98 | 0.58 | 0.87 | 76.68 | 77.46 | 77.07 | 0.54 | 0.85 |
| Pat11 (ETree) | 81.54 | 76.32 | 78.93 | 0.58 | 0.87 | 75.39 | 78.50 | 76.94 | 0.54 | 0.86 |
| Pat13 (ETree) | 79.29 | 80.42 | 79.85 | 0.60 | 0.88 | 74.87 | 81.61 | 78.24 | 0.57 | 0.86 |
| Pat15 (ETree) | 81.59 | 77.10 | 79.35 | 0.59 | 0.88 | 76.68 | 78.24 | 77.46 | 0.55 | 0.86 |
| Pat17 (ETree) | 79.24 | 81.54 | 80.39 | 0.61 | 0.88 | 72.54 | 81.61 | 77.07 | 0.54 | 0.86 |
| Pat19 (SVC) | 80.47 | 79.97 | 80.22 | 0.60 | 0.87 | 75.13 | 78.24 | 76.68 | 0.53 | 0.85 |
| Pat21 (SVC) | 80.92 | 78.84 | 79.88 | 0.60 | 0.87 | 75.39 | 77.46 | 76.42 | 0.53 | 0.85 |
| Pat23 (ETree) | 79.91 | 81.14 | 80.53 | 0.61 | 0.88 | 73.06 | 81.61 | 77.33 | 0.55 | 0.86 |

\* **Sen:** Sensitivity, **Spc:** Specificity, **Acc:** Accuracy, **MCC:** Matthews Correlation Coefficient, **AUROC:** Area Under the Receiver Operating Characteristic curve, **SVC:** Support Vector Classifier, **RF:** Random Forest, **ETree:** Extratree

**Figure 3.**
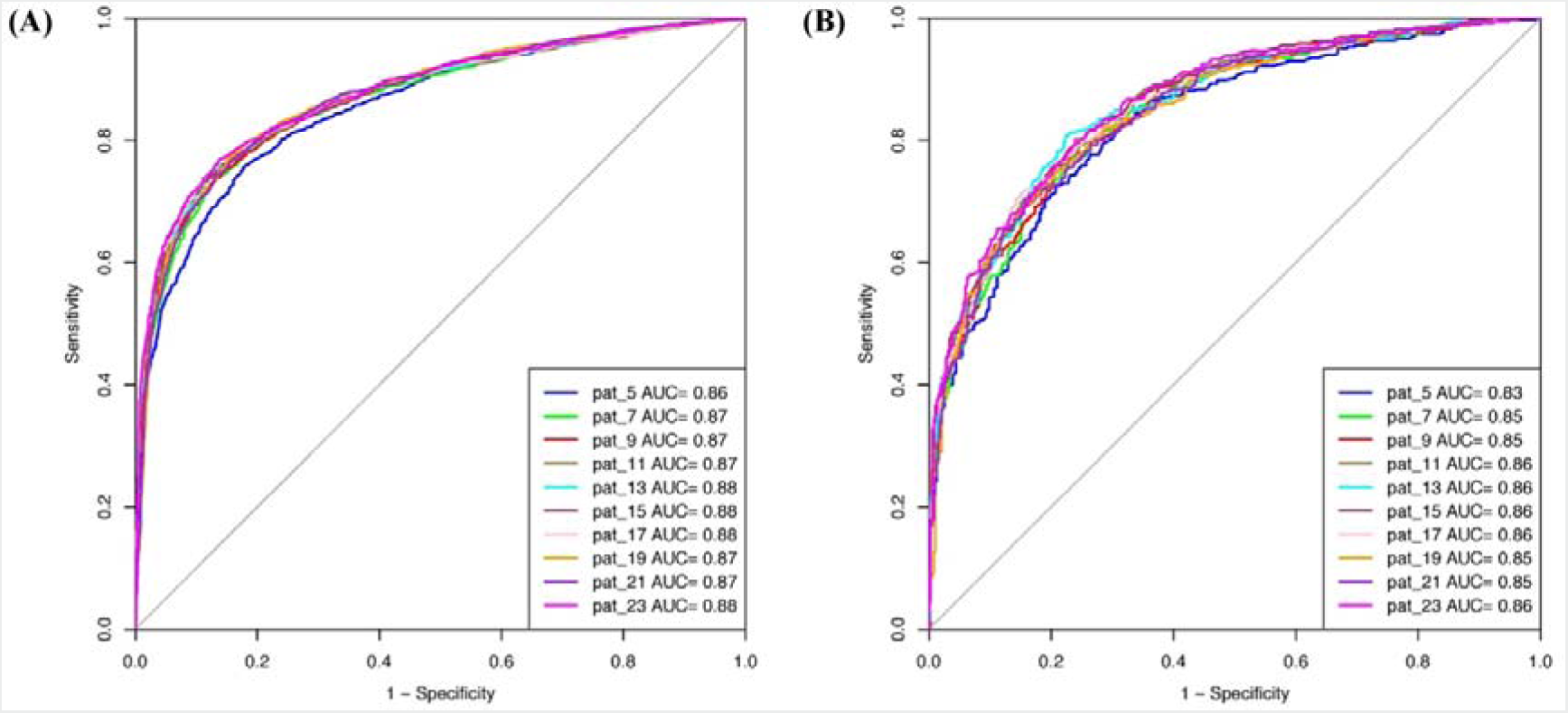
AUROC plots obtained for various window length developed using evolutionary profile on balanced dataset for (a) training dataset and (b) validation dataset.

### Machine learning model performance using hybrid feature

We also developed models on the hybrid feature where we sum up the values of binary profile and the evolutionary information obtained for the residue. Best result obtained for each window length is compiled in the Supplementary Table S21 and plotted AUROC for the training dataset (Supplementary Figure S3 (a)) and validation dataset (Supplementary Figure S3 (b)). Maximum accuracy of 80.58%, MCC of 0.61 and AUROC of 0.89 was obtained by SVC on the training dataset for window size 19. In case of validation dataset, accuracy of 78.50%, MCC of 0.57 and AUROC of 0.87 was obtained. Result for all the window size obtained by different classifiers is provided in the **Supplementary Table S22-S31.**

### Performance of the machine learning models developed on the realistic dataset

Window size 17 was found to be optimum window size as the model developed using PSSM profile performed best among all the models. Therefore, we used this window size for developing prediction models on the realistic dataset using PSSM profile as an input feature. When balanced specificity and sensitivity was considered SVC based model achieved maximum MCC value of 0.32 on training dataset and 0.31 on independent dataset. However, MCC value increases to 0.61 on training dataset and 0.52 on validation dataset when balanced sensitivity and specificity was not taken into account (Table 3). The AUROC achieved on the training dataset and validation dataset was 0.89 and 0.87 respectively **(Figure 4)**.

**Table 3.** The performance of PSSM profile based models developed using different machine learning techniques for window size 17 on realistic dataset.

| Machine Learning<br>Techniques<br>(Parameters) | Main Dataset |  |  |  |  | Validation Dataset |  |  |  |  |
| --- | --- | --- | --- | --- | --- | --- | --- | --- | --- | --- |
|  | Sen | Spc | Acc | MCC | AUROC | Sen | Spc | Acc | MCC | AUROC |
| SVC<sub>MaxMCC#</sub><br>(g=0.01, c=5) | 53.65 | 98.94 | 96.64 | 0.61 | 0.89 | 36.53 | 99.40 | 95.99 | 0.52 | 0.87 |
| SVC<sub>balanced*</sub><br>(g=0.01, c=5) | 81.48 | 80.14 | 80.21 | 0.32 | 0.89 | 77.72 | 79.88 | 79.76 | 0.31 | 0.87 |
| Random Forest<sub>MaxMCC#</sub><br>(Ntree = 700) | 43.27 | 99.44 | 96.59 | 0.58 | 0.86 | 36.53 | 98.97 | 95.58 | 0.48 | 0.86 |
| Random Forest<sub>balanced*</sub><br>(Ntree = 700) | 80.25 | 74.64 | 74.93 | 0.27 | 0.86 | 76.94 | 79.73 | 79.58 | 0.30 | 0.86 |
| Extra Tree<sub>MaxMCC#</sub><br>(Ntree = 1000) | 46.13 | 99.42 | 96.72 | 0.60 | 0.87 | 47.67 | 97.93 | 95.20 | 0.50 | 0.86 |
| Extra Tree<sub>balanced*</sub><br>(Ntree = 1000) | 78.90 | 79.79 | 79.74 | 0.31 | 0.87 | 74.09 | 83.70 | 83.18 | 0.33 | 0.86 |
| Nearest Neighbors<sub>MaxMCC#</sub><br>(neighbors=10,<br>algorithm=auto,<br>weights=distance) | 46.91 | 99.27 | 96.61 | 0.59 | 0.84 | 34.72 | 99.36 | 95.85 | 0.50 | 0.79 |
| Nearest Neighbors<sub>balanced*</sub><br>(neighbors=10,<br>algorithm=auto,<br>weights=distance) | 76.04 | 79.72 | 79.53 | 0.29 | 0.84 | 68.13 | 80.56 | 79.89 | 0.27 | 0.79 |
| Muti Layer<br>Perceptron<sub>MaxMCC#</sub><br>(activation=relu,<br>solver=adam,<br>hidden_layer_sizes=21,<br>max_iter=10) | 37.49 | 98.59 | 95.49 | 0.45 | 0.85 | 34.46 | 98.01 | 94.55 | 0.39 | 0.83 |
| Muti Layer<br>Perceptron<sub>balanced*</sub><br>(activation=relu,<br>solver=adam,<br>hidden_layer_sizes=21,<br>max_iter=10) | 78.84 | 75.84 | 76.00 | 0.27 | 0.85 | 65.28 | 83.26 | 82.28 | 0.28 | 0.83 |
| Ridge Classifier<sub>MaxMCC#</sub><br>(alpha=1) | 38.61 | 97.55 | 94.55 | 0.39 | 0.83 | 37.31 | 96.13 | 92.93 | 0.33 | 0.80 |
| Ridge Classifier<sub>balanced*</sub><br>(alpha=1) | 79.57 | 67.37 | 67.99 | 0.22 | 0.83 | 77.46 | 67.42 | 67.97 | 0.21 | 0.80 |

**Note:** **Sen:** Sensitivity, **Sp:** Specificity, **Acc:** Accuracy, **MCC:** Matthews Correlation Coefficient, **AUROC:** Area Under the Receiver Operating Characteristic curve.
**\*: Results are reported on balanced sensitivity and specificity; #: Results are reported for which maximum MCC was obtained.**

**Figure 4.**
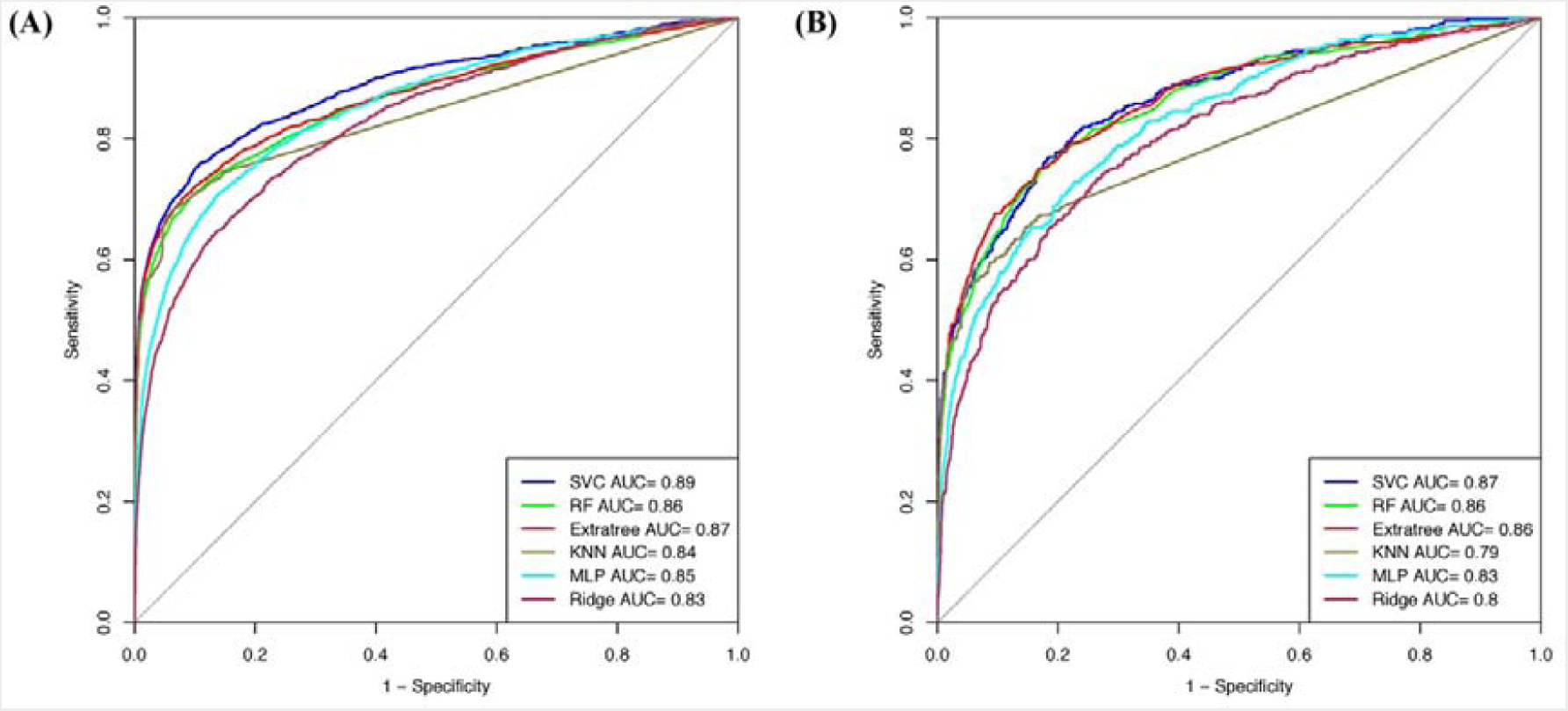
AUROC plots obtained for window length 17 developed using evolutionary profile on realistic dataset for (a) training dataset and (b) validation dataset.

### Performance on additional dataset

We also checked the performance of our best model developed on realistic dataset using PSSM profile and window size 17 on an additional dataset. As shown in Supplementary Table S32, model developed using SVC classifier achieved the best performance with accuracy of 82.27%, and 0.33 MCC with balanced sensitivity and specificity. However the maximum accuracy of 97.14% and MCC of 0.62 was obtained when no balanced sensitivity and specificity were considered. The AUROC obtained was 0.88.

### Implementation of Model in Web server

In order to help biologist for predicting SAM interacting residues, we implemented our best models in a web server “SAMbinder”. The web server consists of several modules such as “Sequence”, “PSSM Profile”, “Peptide Mapping”, “Standalone” and “Download”. These modules have been explained below in detail.

#### (i) Sequence

This module allows users to predict SAM interacting residue in a protein from its primary sequence. A user can submit either single or multiple sequences or upload the sequence file in the FASTA format and can select the desired probability cut-off and machine learning classifier for prediction. The module utilizes the binary profile as an input feature and number of machine learning models has been implemented into it. The classifier provides the prediction score which is normalized in between propensity score 0-9. Residues having the propensity score equal or above the selected cut-off threshold are highlighted in blue colour and remaining residues are highlighted in black colour. Blue colour indicates that the probability of these residues in SAM binding is high in comparison to the residues present in black colour. The result is downloadable in the “csv” file format and will be sent to email also if the user has provided the email.

#### (ii) PSSM Profile

As the name suggests, this module utilizes the PSSM profile as an input feature for predicting SAM interacting residues in a given protein sequence. This feature is better than the binary profile however the only limitation is that it is very computer intensive. Therefore, a user can use this module if the number of the sequences are very few. The output is provided in the same format as Sequence module provides. For doing the prediction for multiple sequences using PSSM profile, we suggest user to use the standalone version of the software.

#### (iii) Peptide Mapping

In this module, we have provided the facility where a user can map the peptide that contains SAM interacting central residue. We pre-computed propensity (between 0-9) of each tri and penta-peptides which contains SAM interacting central residues from known PDB protein structure. The propensity was computed using all SAM interacting protein chains i.e. redundancy was not removed, in order to avoid loss of information. Once a user submits sequence in FASTA format, all the possible segment of selected length is generated and mapped on the protein sequence along with the propensity score. Based on that mapping server predicts whether the peptide segment is SAM interacting or non-interacting. If propensity of residue is equal to greater than the selected threshold, it is known as SAM interacting residue.

### Standalone

Standalone of SAMbinder is Python-based and is available at the Github site. The user can download it from the site https://github.com/raghavagps/sambinder/. SAMbinder standalone version is also implemented in the docker technology. Complete usage of downloading the image and its implementation is provided in the docker manual “GPSRdocker” which can be downloaded for the website https://webs.iiitd.edu.in/gpsrdocker/.

## Discussion

SAM is one of the important essential metabolic cofactor/intermediates which is found in almost every cellular life forms and enzymes. SAM binding proteins are predominant in two major types of folds; (i) Rossman fold and TIM barrel fold and different motifs (Motif I-VI) (Gana *et al*., 2013). SAM binding proteins play a vital role in many metabolic and regulatory pathways in almost all form of living organism and acts as a potential drug target in a number of diseases. In Europe, SAMe is used as drug for treating diseases like liver disorder, depression, fibromyalgia and osteoarthritis. It has also been used as dietary supplements in United States for supporting the bones and joints. Therefore, it is very important to predict the SAM interacting residues in a given protein. We analysed various properties of SAM interacting protein chains such as composition, propensity and physiochemical properties and developed various machine learning models for predicting SAM interacting residue in new protein using number of input features. The models were first developed on the balanced dataset and different window sizes. We observed that model developed using PSSM profile and window size 17 performed best among all the models. Performance of the models was also validated on an independent dataset and an additional dataset. Python-based machine learning package scikit-learn was implemented for developing the prediction models. In order to assist the scientific community, we have created a python-based standalone version of our software and also developed a web server where a user can predict the SAM interacting residues in the target protein. The server can be freely accessible at http://webs.iiitd.edu.in/raghava/sambinder. Complete workflow of SAMbinder is shown in figure 5.

**Figure 5.**
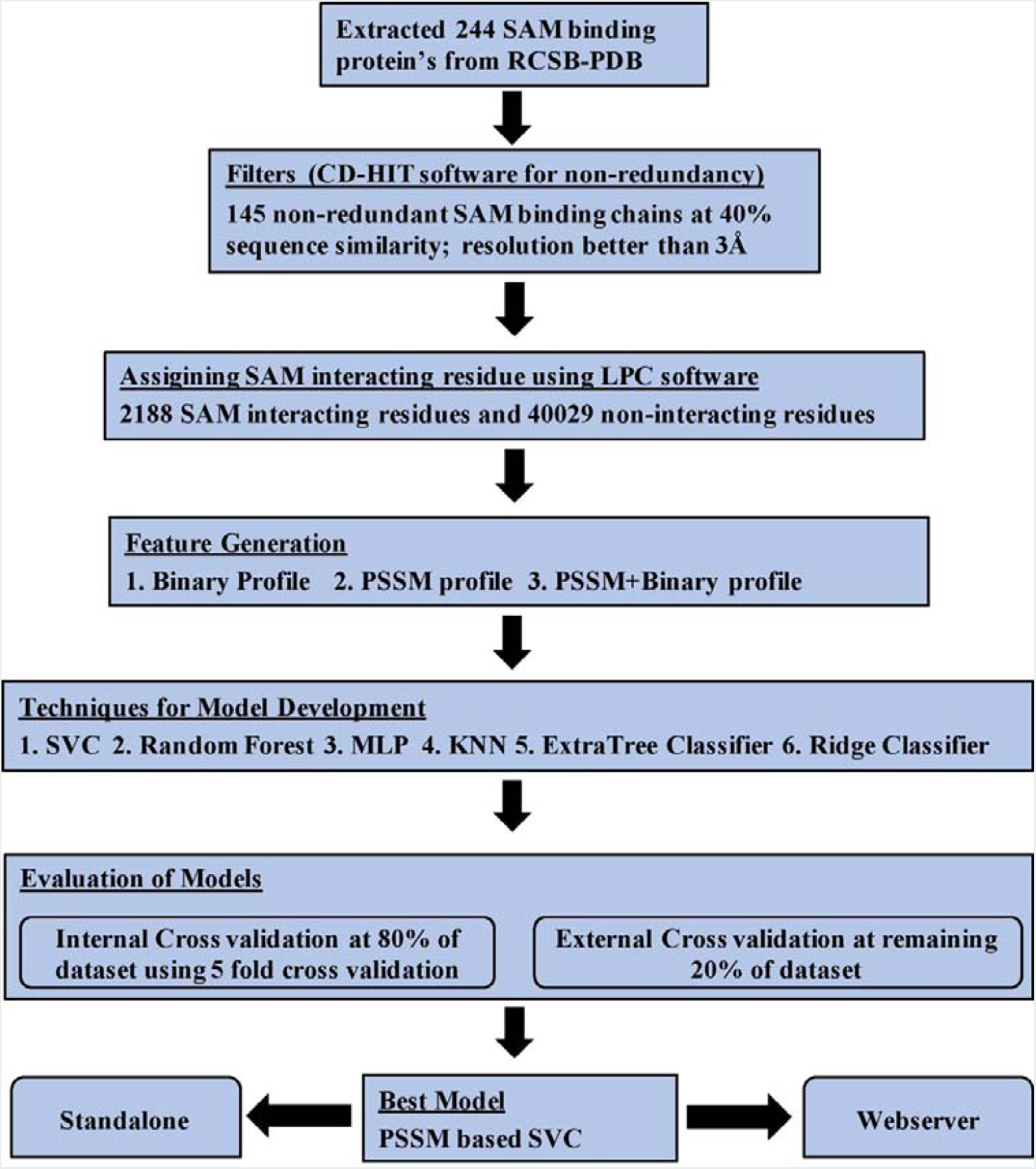
Architecture of SAMbinder.

## Author’s Contribution

PA collected and compiled the datasets. PA performed the experiments. PA and GPSR developed the web interface. PA and GM developed the standalone software. PA, and GPSR analysed the data and prepared the manuscript. GPSR conceived the idea and coordinated the project. All authors read and approved the final paper.

## Acknowledgement

Authors are thankful to J.C. Bose National Fellowship, Department of Science and Technology (DST), Government of India, and DST-INSPIRE for fellowships and the financial support.

