## Supplementary data for "SAMbinder: A web server for predicting SAM binding residues of a protein from its amino acid sequence"

Prof. G.P.S. Raghava,

Head of Department, Department of Computational Biology, Indraprastha Institute of Information Technology, Okhla Phase 3, New Delhi-110020, India.

Phone No: +91-11-26907444

**Supplementary Information**

**Table S1. The performance of different machine learning models developed using amino acid sequence (binary pattern) with window length 5 on balanced dataset.**

| **Machine Learning Techniques**  (**Parameters**) | **Main Dataset** | | | | | **Validation Dataset** | | | | |
| --- | --- | --- | --- | --- | --- | --- | --- | --- | --- | --- |
|  | **Sen** | **Spc** | **Acc** | **MCC** | **AUROC** | **Sen** | **Spc** | **Acc** | **MCC** | **AUROC** |
| SVC  (g=, c=) | 64.98 | 64.31 | 64.65 | 0.29 | 0.70 | 65.54 | 68.65 | 67.10 | 0.34 | 0.72 |
| Random Forest  (Ntree=) | 65.71 | 61.67 | 63.69 | 0.27 | 0.70 | 66.06 | 63.21 | 64.64 | 0.29 | 0.71 |
| ExtraTree Classifier  (Ntree=) | 63.24 | 60.55 | 61.90 | 0.24 | 0.68 | 67.10 | 62.44 | 64.77 | 0.30 | 0.70 |
| K Nearest Neighbor  (n_neighbors=, algorithm=, weights=) | 64.70 | 59.99 | 62.35 | 0.25 | 0.67 | 63.99 | 64.77 | 64.38 | 0.29 | 0.69 |
| Multi Layer Perceptron  (activation=, solver=, hidden layer size=, maximum iteration=10) | 61.22 | 63.47 | 62.35 | 0.25 | 0.68 | 65.54 | 62.95 | 64.25 | 0.29 | 0.70 |
| Ridge Classifier  (alpha=1) | 63.41 | 63.08 | 63.24 | 0.26 | 0.69 | 63.73 | 66.32 | 65.03 | 0.30 | 0.71 |

*** Sen:** Sensitivity, **Spc:** Specificity, **Acc:** Accuracy, **MCC:** Matthews Correlation Coefficient, **AUROC:** Area Under the Receiver Operating Characteristic curve.

**Table S2. The performance of different machine learning models developed using amino acid sequence (binary pattern) with window length 7 on balanced dataset.**

| **Machine Learning Techniques**  (**Parameters**) | **Main Dataset** | | | | | **Validation Dataset** | | | | |
| --- | --- | --- | --- | --- | --- | --- | --- | --- | --- | --- |
|  | **Sen** | **Spc** | **Acc** | **MCC** | **AUROC** | **Sen** | **Spc** | **Acc** | **MCC** | **AUROC** |
| SVC  (g=, c=) | 68.63 | 65.49 | 67.06 | 0.34 | 0.74 | 68.91 | 68.91 | 68.91 | 0.38 | 0.75 |
| Random Forest  (Ntree=) | 69.87 | 64.31 | 67.09 | 0.34 | 0.74 | 69.43 | 66.58 | 68.01 | 0.36 | 0.74 |
| ExtraTree Classifier  (Ntree=) | 67.56 | 64.14 | 65.85 | 0.32 | 0.73 | 65.28 | 66.32 | 65.80 | 0.32 | 0.73 |
| K Nearest Neighbor  (n_neighbors=, algorithm=, weights=) | 67.56 | 62.40 | 64.98 | 0.30 | 0.70 | 65.54 | 68.13 | 66.84 | 0.34 | 0.73 |
| Multi Layer Perceptron  (activation=, solver=, hidden layer size=, maximum iteration=10) | 67.06 | 62.23 | 64.65 | 0.29 | 0.71 | 68.65 | 62.95 | 65.80 | 0.32 | 0.72 |
| Ridge Classifier  (alpha=1) | 64.87 | 65.66 | 65.26 | 0.31 | 0.71 | 66.58 | 66.84 | 66.71 | 0.33 | 0.73 |

*** Sen:** Sensitivity, **Spc:** Specificity, **Acc:** Accuracy, **MCC:** Matthews Correlation Coefficient, **AUROC:** Area Under the Receiver Operating Characteristic curve.

**Table S3. The performance of different machine learning models developed using amino acid sequence (binary pattern) with window length 9 on balanced dataset.**

| **Machine Learning Techniques**  (**Parameters**) | **Main Dataset** | | | | | **Validation Dataset** | | | | |
| --- | --- | --- | --- | --- | --- | --- | --- | --- | --- | --- |
|  | **Sen** | **Spc** | **Acc** | **MCC** | **AUROC** | **Sen** | **Spc** | **Acc** | **MCC** | **AUROC** |
| SVC  (g=, c=) | 67.34 | 66.39 | 66.86 | 0.34 | 0.73 | 69.43 | 69.43 | 69.43 | 0.39 | 0.75 |
| Random Forest  (Ntree=) | 72.05 | 65.66 | 68.86 | 0.38 | 0.76 | 69.95 | 68.65 | 69.30 | 0.39 | 0.76 |
| ExtraTree Classifier  (Ntree=) | 69.81 | 65.99 | 67.90 | 0.36 | 0.75 | 66.06 | 67.88 | 66.97 | 0.34 | 0.75 |
| K Nearest Neighbor  (n_neighbors=, algorithm=, weights=) | 71.10 | 61.95 | 66.53 | 0.33 | 0.73 | 66.06 | 66.32 | 66.19 | 0.32 | 0.71 |
| Multi Layer Perceptron  (activation=, solver=, hidden layer size=, maximum iteration=10) | 66.89 | 65.21 | 66.05 | 0.32 | 0.72 | 70.21 | 64.77 | 67.49 | 0.35 | 0.74 |
| Ridge Classifier  (alpha=1) | 69.30 | 62.18 | 65.74 | 0.32 | 0.72 | 72.54 | 66.06 | 69.30 | 0.39 | 0.75 |

*** Sen:** Sensitivity, **Spc:** Specificity, **Acc:** Accuracy, **MCC:** Matthews Correlation Coefficient, **AUROC:** Area Under the Receiver Operating Characteristic curve.

**Table S4. The performance of different machine learning models developed using amino acid sequence (binary pattern) with window length 11 on balanced dataset.**

| **Machine Learning Techniques**  (**Parameters**) | **Main Dataset** | | | | | **Validation Dataset** | | | | |
| --- | --- | --- | --- | --- | --- | --- | --- | --- | --- | --- |
|  | **Sen** | **Spc** | **Acc** | **MCC** | **AUROC** | **Sen** | **Spc** | **Acc** | **MCC** | **AUROC** |
| SVC  (g=, c=) | 68.52 | 66.44 | 67.48 | 0.35 | 0.74 | 68.91 | 67.88 | 68.39 | 0.37 | 0.76 |
| Random Forest  (Ntree=) | 69.19 | 70.26 | 69.73 | 0.39 | 0.77 | 65.03 | 71.76 | 68.39 | 0.37 | 0.76 |
| ExtraTree Classifier  (Ntree=) | 71.55 | 66.95 | 69.25 | 0.39 | 0.76 | 67.88 | 70.21 | 69.04 | 0.38 | 0.76 |
| K Nearest Neighbor  (n_neighbors=, algorithm=, weights=) | 66.55 | 66.61 | 66.58 | 0.33 | 0.73 | 63.21 | 69.69 | 66.45 | 0.33 | 0.73 |
| Multi Layer Perceptron  (activation=, solver=, hidden layer size=, maximum iteration=10) | 67.90 | 64.09 | 65.99 | 0.32 | 0.72 | 68.91 | 69.69 | 69.30 | 0.39 | 0.76 |
| Ridge Classifier  (alpha=1) | 67.85 | 67.28 | 67.56 | 0.35 | 0.73 | 69.69 | 68.65 | 69.17 | 0.38 | 0.76 |

*** Sen:** Sensitivity, **Spc:** Specificity, **Acc:** Accuracy, **MCC:** Matthews Correlation Coefficient, **AUROC:** Area Under the Receiver Operating Characteristic curve.

**Table S5. The performance of different machine learning models developed using amino acid sequence (binary pattern) with window length 13 on balanced dataset.**

| **Machine Learning Techniques**  (**Parameters**) | **Main Dataset** | | | | | **Validation Dataset** | | | | |
| --- | --- | --- | --- | --- | --- | --- | --- | --- | --- | --- |
|  | **Sen** | **Spc** | **Acc** | **MCC** | **AUROC** | **Sen** | **Spc** | **Acc** | **MCC** | **AUROC** |
| SVC  (g=, c=) | 68.41 | 67.68 | 68.04 | 0.36 | 0.74 | 69.43 | 69.43 | 69.43 | 0.39 | 0.76 |
| Random Forest  (Ntree=) | 73.06 | 66.05 | 69.56 | 0.39 | 0.77 | 70.98 | 65.03 | 68.01 | 0.36 | 0.77 |
| ExtraTree Classifier  (Ntree=) | 71.66 | 67.17 | 69.42 | 0.39 | 0.77 | 68.91 | 69.17 | 69.04 | 0.38 | 0.76 |
| K Nearest Neighbor  (n_neighbors=, algorithm=, weights=) | 67.45 | 67.51 | 67.48 | 0.35 | 0.75 | 61.66 | 68.39 | 65.03 | 0.30 | 0.73 |
| Multi Layer Perceptron  (activation=, solver=, hidden layer size=, maximum iteration=10) | 70.71 | 64.53 | 67.62 | 0.35 | 0.74 | 72.54 | 66.32 | 69.43 | 0.39 | 0.76 |
| Ridge Classifier  (alpha=1) | 67.62 | 67.17 | 67.40 | 0.35 | 0.73 | 69.95 | 68.91 | 69.43 | 0.39 | 0.77 |

*** Sen:** Sensitivity, **Spc:** Specificity, **Acc:** Accuracy, **MCC:** Matthews Correlation Coefficient, **AUROC:** Area Under the Receiver Operating Characteristic curve.

**Table S6. The performance of different machine learning models developed using amino acid sequence (binary pattern) with window length 15 on balanced dataset.**

| **Machine Learning Techniques**  (**Parameters**) | **Main Dataset** | | | | | **Validation Dataset** | | | | |
| --- | --- | --- | --- | --- | --- | --- | --- | --- | --- | --- |
|  | **Sen** | **Spc** | **Acc** | **MCC** | **AUROC** | **Sen** | **Spc** | **Acc** | **MCC** | **AUROC** |
| SVC  (g=, c=) | 69.53 | 66.89 | 68.21 | 0.36 | 0.74 | 69.17 | 70.73 | 69.95 | 0.40 | 0.77 |
| Random Forest  (Ntree=) | 69.58 | 70.71 | 70.15 | 0.40 | 0.78 | 66.06 | 72.02 | 69.04 | 0.38 | 0.78 |
| ExtraTree Classifier  (Ntree=) | 72.33 | 68.13 | 70.23 | 0.40 | 0.77 | 69.95 | 68.91 | 69.43 | 0.39 | 0.77 |
| K Nearest Neighbor  (n_neighbors=, algorithm=, weights=) | 69.47 | 65.49 | 67.48 | 0.35 | 0.75 | 63.47 | 69.95 | 66.71 | 0.33 | 0.73 |
| Multi Layer Perceptron  (activation=, solver=, hidden layer size=, maximum iteration=10) | 67.85 | 66.27 | 67.06 | 0.34 | 0.74 | 71.50 | 72.02 | 71.76 | 0.44 | 0.77 |
| Ridge Classifier  (alpha=1) | 67.62 | 66.95 | 67.28 | 0.35 | 0.73 | 69.69 | 69.69 | 69.69 | 0.39 | 0.77 |

*** Sen:** Sensitivity, **Spc:** Specificity, **Acc:** Accuracy, **MCC:** Matthews Correlation Coefficient, **AUROC:** Area Under the Receiver Operating Characteristic curve.

**Table S7. The performance of different machine learning models developed using amino acid sequence (binary pattern) with window length 17 on balanced dataset.**

| **Machine Learning Techniques**  (**Parameters**) | **Main Dataset** | | | | | **Validation Dataset** | | | | |
| --- | --- | --- | --- | --- | --- | --- | --- | --- | --- | --- |
|  | **Sen** | **Spc** | **Acc** | **MCC** | **AUROC** | **Sen** | **Spc** | **Acc** | **MCC** | **AUROC** |
| SVC  (g=, c=) | 68.91 | 68.29 | 68.60 | 0.37 | 0.74 | 68.13 | 70.73 | 69.43 | 0.39 | 0.78 |
| Random Forest  (Ntree=) | 70.37 | 71.10 | 70.74 | 0.41 | 0.78 | 67.36 | 71.50 | 69.43 | 0.39 | 0.78 |
| ExtraTree Classifier  (Ntree=) | 73.34 | 67.40 | 70.37 | 0.41 | 0.78 | 71.24 | 66.84 | 69.04 | 0.38 | 0.77 |
| K Nearest Neighbor  (n_neighbors=, algorithm=, weights=) | 64.53 | 72.84 | 68.69 | 0.38 | 0.75 | 60.62 | 78.24 | 69.43 | 0.39 | 0.74 |
| Multi Layer Perceptron  (activation=, solver=, hidden layer size=, maximum iteration=10) | 69.92 | 64.59 | 67.26 | 0.35 | 0.74 | 72.28 | 69.43 | 70.85 | 0.42 | 0.78 |
| Ridge Classifier  (alpha=1) | 68.74 | 66.72 | 67.73 | 0.35 | 0.73 | 71.24 | 70.21 | 70.73 | 0.41 | 0.77 |

*** Sen:** Sensitivity, **Spc:** Specificity, **Acc:** Accuracy, **MCC:** Matthews Correlation Coefficient, **AUROC:** Area Under the Receiver Operating Characteristic curve.

**Table S8. The performance of different machine learning models developed using amino acid sequence (binary pattern) with window length 19 on balanced dataset.**

| **Machine Learning Techniques**  (**Parameters**) | **Main Dataset** | | | | | **Validation Dataset** | | | | |
| --- | --- | --- | --- | --- | --- | --- | --- | --- | --- | --- |
|  | **Sen** | **Spc** | **Acc** | **MCC** | **AUROC** | **Sen** | **Spc** | **Acc** | **MCC** | **AUROC** |
| SVC  (g=, c=) | 69.64 | 66.61 | 68.13 | 0.36 | 0.75 | 70.73 | 70.21 | 70.47 | 0.41 | 0.77 |
| Random Forest  (Ntree=) | 70.54 | 71.32 | 70.93 | 0.42 | 0.78 | 67.36 | 73.83 | 70.60 | 0.41 | 0.79 |
| ExtraTree Classifier  (Ntree=) | 74.69 | 67.56 | 71.13 | 0.42 | 0.78 | 70.47 | 67.10 | 68.78 | 0.38 | 0.78 |
| K Nearest Neighbor  (n_neighbors=, algorithm=, weights=) | 66.67 | 74.58 | 70.62 | 0.41 | 0.77 | 61.14 | 72.54 | 66.84 | 0.34 | 0.74 |
| Multi Layer Perceptron  (activation=, solver=, hidden layer size=, maximum iteration=10) | 68.35 | 67.62 | 67.99 | 0.36 | 0.74 | 69.17 | 70.47 | 69.82 | 0.40 | 0.76 |
| Ridge Classifier  (alpha=1) | 68.07 | 66.05 | 67.06 | 0.34 | 0.73 | 70.98 | 70.73 | 70.85 | 0.42 | 0.77 |

*** Sen:** Sensitivity, **Spc:** Specificity, **Acc:** Accuracy, **MCC:** Matthews Correlation Coefficient, **AUROC:** Area Under the Receiver Operating Characteristic curve.

**Table S9. The performance of different machine learning models developed using amino acid sequence (binary pattern) with window length 21 on balanced dataset.**

| **Machine Learning Techniques**  (**Parameters**) | **Main Dataset** | | | | | **Validation Dataset** | | | | |
| --- | --- | --- | --- | --- | --- | --- | --- | --- | --- | --- |
|  | **Sen** | **Spc** | **Acc** | **MCC** | **AUROC** | **Sen** | **Spc** | **Acc** | **MCC** | **AUROC** |
| SVC  (g=, c=) | 68.74 | 68.35 | 68.35 | 0.37 | 0.75 | 68.91 | 71.76 | 70.34 | 0.41 | 0.77 |
| Random Forest  (Ntree=) | 70.37 | 71.21 | 70.79 | 0.42 | 0.78 | 67.62 | 74.09 | 70.85 | 0.42 | 0.79 |
| ExtraTree Classifier  (Ntree=) | 73.96 | 67.34 | 70.65 | 0.41 | 0.78 | 70.73 | 67.36 | 69.04 | 0.38 | 0.78 |
| K Nearest Neighbor  (n_neighbors=, algorithm=, weights=) | 69.36 | 66.16 | 67.76 | 0.36 | 0.75 | 64.77 | 69.69 | 67.23 | 0.34 | 0.74 |
| Multi Layer Perceptron  (activation=, solver=, hidden layer size=, maximum iteration=10) | 70.48 | 64.87 | 67.68 | 0.35 | 0.75 | 70.47 | 71.50 | 70.98 | 0.42 | 0.77 |
| Ridge Classifier  (alpha=1) | 67.79 | 66.67 | 67.23 | 0.34 | 0.73 | 70.21 | 69.95 | 70.08 | 0.40 | 0.76 |

*** Sen:** Sensitivity, **Spc:** Specificity, **Acc:** Accuracy, **MCC:** Matthews Correlation Coefficient, **AUROC:** Area Under the Receiver Operating Characteristic curve.

**Table S10. The performance of different machine learning models developed using amino acid sequence (binary pattern) with window length 23 on balanced dataset.**

| **Machine Learning Techniques**  (**Parameters**) | **Main Dataset** | | | | | **Validation Dataset** | | | | |
| --- | --- | --- | --- | --- | --- | --- | --- | --- | --- | --- |
|  | **Sen** | **Spc** | **Acc** | **MCC** | **AUROC** | **Sen** | **Spc** | **Acc** | **MCC** | **AUROC** |
| SVC  (g=, c=) | 68.52 | 69.08 | 68.80 | 0.38 | 0.75 | 69.43 | 70.73 | 70.08 | 0.40 | 0.77 |
| Random Forest  (Ntree=) | 70.76 | 70.99 | 70.88 | 0.42 | 0.78 | 68.39 | 71.76 | 70.08 | 0.40 | 0.79 |
| ExtraTree Classifier  (Ntree=) | 74.19 | 66.44 | 70.31 | 0.41 | 0.78 | 72.28 | 66.58 | 69.43 | 0.39 | 0.78 |
| K Nearest Neighbor  (n_neighbors=, algorithm=, weights=) | 69.30 | 67.79 | 68.55 | 0.37 | 0.76 | 67.88 | 69.43 | 68.85 | 0.37 | 0.75 |
| Multi Layer Perceptron  (activation=, solver=, hidden layer size=, maximum iteration=10) | 68.80 | 67.90 | 68.35 | 0.37 | 0.75 | 69.43 | 70.47 | 69.95 | 0.40 | 0.76 |
| Ridge Classifier  (alpha=1) | 67.17 | 67.12 | 67.14 | 0.34 | 0.73 | 69.43 | 68.13 | 68.78 | 0.38 | 0.76 |

*** Sen:** Sensitivity, **Spc:** Specificity, **Acc:** Accuracy, **MCC:** Matthews Correlation Coefficient, **AUROC:** Area Under the Receiver Operating Characteristic curve.

**Table S11. The performance of different machine learning models developed using PSSM profile with window length 5 on balanced dataset.**

| **Machine Learning Techniques**  (**Parameters**) | **Main Dataset** | | | | | **Validation Dataset** | | | | |
| --- | --- | --- | --- | --- | --- | --- | --- | --- | --- | --- |
|  | **Sen** | **Spc** | **Acc** | **MCC** | **AUROC** | **Sen** | **Spc** | **Acc** | **MCC** | **AUROC** |
| SVC  (g=, c=) | 75.42 | 74.30 | 74.86 | 0.50 | 0.83 | 73.88 | 68.91 | 71.37 | 0.43 | 0.79 |
| Random Forest  (Ntree=) | 78.40 | 77.50 | 77.95 | 0.56 | 0.86 | 74.35 | 76.94 | 75.65 | 0.51 | 0.83 |
| ExtraTree Classifier  (Ntree=) | 79.18 | 75.76 | 77.47 | 0.55 | 0.86 | 73.83 | 76.17 | 75.00 | 0.50 | 0.83 |
| K Nearest Neighbor  (n_neighbors=, algorithm=, weights=) | 76.99 | 71.16 | 74.07 | 0.48 | 0.83 | 69.69 | 72.28 | 70.98 | 0.42 | 0.78 |
| Multi Layer Perceptron  (activation=, solver=, hidden layer size=, maximum iteration=10) | 72.56 | 72.78 | 72.67 | 0.45 | 0.80 | 69.17 | 67.36 | 68.26 | 0.37 | 0.75 |
| Ridge Classifier  (alpha=1) | 72.90 | 70.43 | 71.66 | 0.43 | 0.79 | 73.58 | 63.73 | 68.65 | 0.37 | 0.75 |

*** Sen:** Sensitivity, **Spc:** Specificity, **Acc:** Accuracy, **MCC:** Matthews Correlation Coefficient, **AUROC:** Area Under the Receiver Operating Characteristic curve.

**Table S12. The performance of different machine learning models developed using PSSM profile with window length 7 on balanced dataset.**

| **Machine Learning Techniques**  (**Parameters**) | **Main Dataset** | | | | | **Validation Dataset** | | | | |
| --- | --- | --- | --- | --- | --- | --- | --- | --- | --- | --- |
|  | **Sen** | **Spc** | **Acc** | **MCC** | **AUROC** | **Sen** | **Spc** | **Acc** | **MCC** | **AUROC** |
| SVC  (g=, c=) | 78.23 | 78.17 | 78.20 | 0.56 | 0.86 | 74.87 | 74.09 | 74.48 | 0.49 | 0.82 |
| Random Forest  (Ntree=) | 81.20 | 76.54 | 78.87 | 0.58 | 0.86 | 75.13 | 79.79 | 77.46 | 0.55 | 0.84 |
| ExtraTree Classifier  (Ntree=) | 79.85 | 79.12 | 79.49 | 0.59 | 0.87 | 73.32 | 79.53 | 76.42 | 0.53 | 0.85 |
| K Nearest Neighbor  (n_neighbors=, algorithm=, weights=) | 76.26 | 72.95 | 74.61 | 0.49 | 0.84 | 69.43 | 72.02 | 70.73 | 0.41 | 0.77 |
| Multi Layer Perceptron  (activation=, solver=, hidden layer size=, maximum iteration=10) | 73.51 | 70.71 | 72.11 | 0.44 | 0.79 | 67.10 | 71.50 | 69.30 | 0.39 | 0.76 |
| Ridge Classifier  (alpha=1) | 75.20 | 73.06 | 74.13 | 0.48 | 0.81 | 73.83 | 67.36 | 70.60 | 0.41 | 0.78 |

*** Sen:** Sensitivity, **Spc:** Specificity, **Acc:** Accuracy, **MCC:** Matthews Correlation Coefficient, **AUROC:** Area Under the Receiver Operating Characteristic curve.

**Table S13. The performance of different machine learning models developed using PSSM profile with window length 9 on balanced dataset.**

| **Machine Learning Techniques**  (**Parameters**) | **Main Dataset** | | | | | **Validation Dataset** | | | | |
| --- | --- | --- | --- | --- | --- | --- | --- | --- | --- | --- |
|  | **Sen** | **Spc** | **Acc** | **MCC** | **AUROC** | **Sen** | **Spc** | **Acc** | **MCC** | **AUROC** |
| SVC  (g=, c=) | 78.90 | 78.90 | 78.90 | 0.58 | 0.86 | 73.32 | 74.35 | 73.83 | 0.48 | 0.82 |
| Random Forest  (Ntree=) | 79.85 | 78.06 | 78.96 | 0.58 | 0.87 | 73.83 | 78.24 | 76.04 | 0.52 | 0.85 |
| ExtraTree Classifier  (Ntree=) | 81.48 | 76.49 | 78.98 | 0.58 | 0.87 | 76.68 | 77.46 | 77.07 | 0.54 | 0.85 |
| K Nearest Neighbor  (n_neighbors=, algorithm=, weights=) | 75.48 | 75.08 | 75.28 | 0.51 | 0.84 | 69.17 | 74.35 | 71.76 | 0.44 | 0.78 |
| Multi Layer Perceptron  (activation=, solver=, hidden layer size=, maximum iteration=10) | 75.36 | 74.13 | 74.75 | 0.49 | 0.82 | 65.03 | 72.80 | 68.91 | 0.38 | 0.77 |
| Ridge Classifier  (alpha=1) | 74.86 | 73.63 | 74.24 | 0.48 | 0.81 | 73.32 | 66.06 | 69.69 | 0.39 | 0.77 |

*** Sen:** Sensitivity, **Spc:** Specificity, **Acc:** Accuracy, **MCC:** Matthews Correlation Coefficient, **AUROC:** Area Under the Receiver Operating Characteristic curve.

**Table S14. The performance of different machine learning models developed using PSSM profile with window length 11 on balanced dataset.**

| **Machine Learning Techniques**  (**Parameters**) | **Main Dataset** | | | | | **Validation Dataset** | | | | |
| --- | --- | --- | --- | --- | --- | --- | --- | --- | --- | --- |
|  | **Sen** | **Spc** | **Acc** | **MCC** | **AUROC** | **Sen** | **Spc** | **Acc** | **MCC** | **AUROC** |
| SVC  (g=, c=) | 80.02 | 79.07 | 79.55 | 0.59 | 0.87 | 73.58 | 77.72 | 75.65 | 0.51 | 0.83 |
| Random Forest  (Ntree=) | 79.85 | 78.34 | 79.10 | 0.58 | 0.87 | 73.32 | 79.02 | 76.17 | 0.52 | 0.85 |
| ExtraTree Classifier  (Ntree=) | 81.54 | 76.32 | 78.93 | 0.58 | 0.87 | 75.39 | 78.50 | 76.94 | 0.54 | 0.86 |
| K Nearest Neighbor  (n_neighbors=, algorithm=, weights=) | 73.46 | 80.98 | 77.22 | 0.55 | 0.83 | 66.32 | 82.90 | 74.61 | 0.50 | 0.81 |
| Multi Layer Perceptron  (activation=, solver=, hidden layer size=, maximum iteration=10) | 75.59 | 75.03 | 75.31 | 0.51 | 0.83 | 73.83 | 69.17 | 71.50 | 0.43 | 0.79 |
| Ridge Classifier  (alpha=1) | 75.20 | 73.96 | 74.58 | 0.49 | 0.82 | 71.76 | 68.39 | 70.08 | 0.40 | 0.78 |

*** Sen:** Sensitivity, **Spc:** Specificity, **Acc:** Accuracy, **MCC:** Matthews Correlation Coefficient, **AUROC:** Area Under the Receiver Operating Characteristic curve.

**Table S15. The performance of different machine learning models developed using PSSM profile with window length 13 on balanced dataset.**

| **Machine Learning Techniques**  (**Parameters**) | **Main Dataset** | | | | | **Validation Dataset** | | | | |
| --- | --- | --- | --- | --- | --- | --- | --- | --- | --- | --- |
|  | **Sen** | **Spc** | **Acc** | **MCC** | **AUROC** | **Sen** | **Spc** | **Acc** | **MCC** | **AUROC** |
| SVC  (g=, c=) | 80.25 | 78.84 | 79.55 | 0.59 | 0.87 | 75.39 | 76.42 | 75.91 | 0.52 | 0.84 |
| Random Forest  (Ntree=) | 80.13 | 78.23 | 79.18 | 0.58 | 0.87 | 74.61 | 79.79 | 77.20 | 0.54 | 0.86 |
| ExtraTree Classifier  (Ntree=) | 79.29 | 80.42 | 79.85 | 0.60 | 0.88 | 74.87 | 81.61 | 78.24 | 0.57 | 0.86 |
| K Nearest Neighbor  (n_neighbors=, algorithm=, weights=) | 75.53 | 76.60 | 76.07 | 0.52 | 0.83 | 69.17 | 77.98 | 73.58 | 0.47 | 0.80 |
| Multi Layer Perceptron  (activation=, solver=, hidden layer size=, maximum iteration=10) | 75.87 | 73.23 | 74.55 | 0.49 | 0.82 | 77.46 | 62.69 | 70.08 | 0.41 | 0.78 |
| Ridge Classifier  (alpha=1) | 74.92 | 74.24 | 74.58 | 0.49 | 0.82 | 70.73 | 69.17 | 69.95 | 0.40 | 0.78 |

*** Sen:** Sensitivity, **Spc:** Specificity, **Acc:** Accuracy, **MCC:** Matthews Correlation Coefficient, **AUROC:** Area Under the Receiver Operating Characteristic curve.

**Table S16. The performance of different machine learning models developed using PSSM profile with window length 15 on balanced dataset.**

| **Machine Learning Techniques**  (**Parameters**) | **Main Dataset** | | | | | **Validation Dataset** | | | | |
| --- | --- | --- | --- | --- | --- | --- | --- | --- | --- | --- |
|  | **Sen** | **Spc** | **Acc** | **MCC** | **AUROC** | **Sen** | **Spc** | **Acc** | **MCC** | **AUROC** |
| SVC  (g=, c=) | 79.80 | 78.90 | 79.35 | 0.59 | 0.87 | 77.20 | 75.39 | 76.30 | 0.53 | 0.85 |
| Random Forest  (Ntree=) | 80.75 | 78.73 | 79.74 | 0.59 | 0.87 | 73.83 | 79.79 | 76.81 | 0.54 | 0.86 |
| ExtraTree Classifier  (Ntree=) | 81.59 | 77.10 | 79.35 | 0.59 | 0.88 | 76.68 | 78.24 | 77.46 | 0.55 | 0.86 |
| K Nearest Neighbor  (n_neighbors=, algorithm=, weights=) | 78.68 | 69.42 | 74.05 | 0.48 | 0.83 | 71.24 | 70.98 | 71.11 | 0.42 | 0.81 |
| Multi Layer Perceptron  (activation=, solver=, hidden layer size=, maximum iteration=10) | 75.81 | 72.84 | 74.33 | 0.49 | 0.83 | 69.69 | 75.13 | 72.41 | 0.45 | 0.80 |
| Ridge Classifier  (alpha=1) | 74.35 | 74.64 | 74.49 | 0.49 | 0.82 | 69.17 | 70.21 | 69.69 | 0.39 | 0.78 |

*** Sen:** Sensitivity, **Spc:** Specificity, **Acc:** Accuracy, **MCC:** Matthews Correlation Coefficient, **AUROC:** Area Under the Receiver Operating Characteristic curve.

**Table S17. The performance of different machine learning models developed using PSSM profile with window length 17 on balanced dataset.**

| **Machine Learning Techniques**  (**Parameters**) | **Main Dataset** | | | | | **Validation Dataset** | | | | |
| --- | --- | --- | --- | --- | --- | --- | --- | --- | --- | --- |
|  | **Sen** | **Spc** | **Acc** | **MCC** | **AUROC** | **Sen** | **Spc** | **Acc** | **MCC** | **AUROC** |
| SVC  (g=, c=) | 80.36 | 79.07 | 79.71 | 0.59 | 0.87 | 75.65 | 78.76 | 77.20 | 0.54 | 0.85 |
| Random Forest  (Ntree=) | 80.36 | 78.00 | 79.18 | 0.58 | 0.87 | 73.83 | 78.76 | 76.30 | 0.53 | 0.85 |
| ExtraTree Classifier  (Ntree=) | 79.24 | 81.54 | 80.39 | 0.61 | 0.88 | 72.54 | 81.61 | 77.07 | 0.54 | 0.86 |
| K Nearest Neighbor  (n_neighbors=, algorithm=, weights=) | 74.30 | 73.79 | 74.05 | 0.48 | 0.83 | 69.43 | 76.94 | 73.19 | 0.47 | 0.80 |
| Multi Layer Perceptron  (activation=, solver=, hidden layer size=, maximum iteration=10) | 75.31 | 74.75 | 75.03 | 0.50 | 0.83 | 71.50 | 76.42 | 73.96 | 0.48 | 0.80 |
| Ridge Classifier  (alpha=1) | 75.14 | 74.19 | 74.66 | 0.49 | 0.82 | 68.39 | 71.76 | 70.08 | 0.40 | 0.78 |

*** Sen:** Sensitivity, **Spc:** Specificity, **Acc:** Accuracy, **MCC:** Matthews Correlation Coefficient, **AUROC:** Area Under the Receiver Operating Characteristic curve.

**Table S18. The performance of different machine learning models developed using PSSM profile with window length 19 on balanced dataset.**

| **Machine Learning Techniques**  (**Parameters**) | **Main Dataset** | | | | | **Validation Dataset** | | | | |
| --- | --- | --- | --- | --- | --- | --- | --- | --- | --- | --- |
|  | **Sen** | **Spc** | **Acc** | **MCC** | **AUROC** | **Sen** | **Spc** | **Acc** | **MCC** | **AUROC** |
| SVC  (g=, c=) | 80.47 | 79.97 | 80.22 | 0.60 | 0.87 | 75.13 | 78.24 | 76.68 | 0.53 | 0.85 |
| Random Forest  (Ntree=) | 80.81 | 78.06 | 79.43 | 0.59 | 0.87 | 73.32 | 78.76 | 76.04 | 0.52 | 0.85 |
| ExtraTree Classifier  (Ntree=) | 82.21 | 76.04 | 79.12 | 0.58 | 0.88 | 76.68 | 78.24 | 77.46 | 0.55 | 0.86 |
| K Nearest Neighbor  (n_neighbors=, algorithm=, weights=) | 74.13 | 76.54 | 75.34 | 0.51 | 0.83 | 69.43 | 79.79 | 74.61 | 0.49 | 0.81 |
| Multi Layer Perceptron  (activation=, solver=, hidden layer size=, maximum iteration=10) | 76.60 | 74.13 | 75.36 | 0.51 | 0.83 | 69.95 | 77.20 | 73.58 | 0.47 | 0.82 |
| Ridge Classifier  (alpha=1) | 75.25 | 74.69 | 74.97 | 0.50 | 0.82 | 69.69 | 71.24 | 70.47 | 0.41 | 0.79 |

*** Sen:** Sensitivity, **Spc:** Specificity, **Acc:** Accuracy, **MCC:** Matthews Correlation Coefficient, **AUROC:** Area Under the Receiver Operating Characteristic curve.

**Table S19. The performance of different machine learning models developed using PSSM profile with window length 21 on balanced dataset.**

| **Machine Learning Techniques**  (**Parameters**) | **Main Dataset** | | | | | **Validation Dataset** | | | | |
| --- | --- | --- | --- | --- | --- | --- | --- | --- | --- | --- |
|  | **Sen** | **Spc** | **Acc** | **MCC** | **AUROC** | **Sen** | **Spc** | **Acc** | **MCC** | **AUROC** |
| SVC  (g=, c=) | 80.92 | 78.84 | 79.88 | 0.60 | 0.87 | 75.39 | 77.46 | 76.42 | 0.53 | 0.85 |
| Random Forest  (Ntree=) | 80.81 | 77.89 | 79.35 | 0.59 | 0.87 | 73.83 | 78.50 | 76.17 | 0.52 | 0.86 |
| ExtraTree Classifier  (Ntree=) | 79.85 | 81.59 | 80.72 | 0.61 | 0.88 | 74.87 | 81.87 | 78.37 | 0.57 | 0.86 |
| K Nearest Neighbor  (n_neighbors=, algorithm=, weights=) | 73.12 | 78.96 | 76.04 | 0.52 | 0.83 | 65.54 | 83.94 | 74.74 | 0.50 | 0.81 |
| Multi Layer Perceptron  (activation=, solver=, hidden layer size=, maximum iteration=10) | 76.43 | 73.96 | 75.20 | 0.50 | 0.84 | 67.62 | 80.05 | 73.83 | 0.48 | 0.82 |
| Ridge Classifier  (alpha=1) | 75.48 | 73.91 | 74.69 | 0.49 | 0.82 | 70.98 | 73.58 | 72.28 | 0.45 | 0.79 |

*** Sen:** Sensitivity, **Spc:** Specificity, **Acc:** Accuracy, **MCC:** Matthews Correlation Coefficient, **AUROC:** Area Under the Receiver Operating Characteristic curve.

**Table S20. The performance of different machine learning models developed using PSSM profile with window length 23 on balanced dataset.**

| **Machine Learning Techniques**  (**Parameters**) | **Main Dataset** | | | | | **Validation Dataset** | | | | |
| --- | --- | --- | --- | --- | --- | --- | --- | --- | --- | --- |
|  | **Sen** | **Spc** | **Acc** | **MCC** | **AUROC** | **Sen** | **Spc** | **Acc** | **MCC** | **AUROC** |
| SVC  (g=, c=) | 82.32 | 75.53 | 78.93 | 0.58 | 0.87 | 77.98 | 75.39 | 76.68 | 0.53 | 0.85 |
| Random Forest  (Ntree=) | 80.36 | 77.78 | 79.07 | 0.58 | 0.87 | 75.13 | 76.94 | 76.04 | 0.52 | 0.85 |
| ExtraTree Classifier  (Ntree=) | 79.91 | 81.14 | 80.53 | 0.61 | 0.88 | 73.06 | 81.61 | 77.33 | 0.55 | 0.86 |
| K Nearest Neighbor  (n_neighbors=, algorithm=, weights=) | 77.78 | 71.16 | 74.47 | 0.49 | 0.83 | 68.13 | 74.09 | 71.11 | 0.42 | 0.80 |
| Multi Layer Perceptron  (activation=, solver=, hidden layer size=, maximum iteration=10) | 75.93 | 74.92 | 75.42 | 0.51 | 0.84 | 74.61 | 77.20 | 75.91 | 0.52 | 0.83 |
| Ridge Classifier  (alpha=1) | 75.03 | 74.13 | 74.58 | 0.49 | 0.82 | 73.32 | 74.61 | 73.96 | 0.48 | 0.80 |

*** Sen:** Sensitivity, **Spc:** Specificity, **Acc:** Accuracy, **MCC:** Matthews Correlation Coefficient, **AUROC:** Area Under the Receiver Operating Characteristic curve.

**Table S21. The performance of best machine learning model developed using hybrid feature for individual window size on balanced dataset.**

|  | **Training Dataset** | | | | | **Validation Dataset** | | | | |
| --- | --- | --- | --- | --- | --- | --- | --- | --- | --- | --- |
|  | **Sen** | **Spc** | **Acc** | **MCC** | **AUROC** | **Sen** | **Spc** | **Acc** | **MCC** | **AUROC** |
| Pat5 (ETree) | 80.64 | 76.09 | 78.37 | 0.57 | 0.86 | 75.13 | 75.39 | 75.26 | 0.51 | 0.84 |
| Pat7 (ETree) | 79.91 | 78.62 | 79.26 | 0.59 | 0.87 | 73.83 | 79.79 | 76.81 | 0.54 | 0.85 |
| Pat9 (ETree) | 79.80 | 79.24 | 79.52 | 0.59 | 0.88 | 73.58 | 81.09 | 77.33 | 0.55 | 0.86 |
| Pat11 (ETree) | 82.49 | 76.37 | 79.43 | 0.59 | 0.88 | 77.72 | 80.05 | 78.89 | 0.58 | 0.87 |
| Pat13 (SVC) | 81.82 | 78.00 | 79.91 | 0.60 | 0.88 | 77.20 | 79.02 | 78.11 | 0.56 | 0.86 |
| Pat15 (SVC) | 81.37 | 80.02 | 80.70 | 0.61 | 0.88 | 77.20 | 78.50 | 77.85 | 0.56 | 0.86 |
| Pat17 (SVC) | 81.54 | 79.91 | 80.72 | 0.61 | 0.88 | 78.76 | 77.72 | 78.24 | 0.56 | 0.87 |
| Pat19 (SVC) | 83.50 | 77.67 | 80.58 | 0.61 | 0.89 | 81.09 | 75.91 | 78.50 | 0.57 | 0.87 |
| Pat21 (ETree) | 80.08 | 80.58 | 80.33 | 0.61 | 0.88 | 75.91 | 81.87 | 78.89 | 0.58 | 0.87 |
| Pat23 (ETree) | 80.86 | 79.35 | 80.11 | 0.60 | 0.88 | 74.87 | 82.38 | 78.63 | 0.57 | 0.87 |

*** Sen:** Sensitivity, **Spc:** Specificity, **Acc:** Accuracy, **MCC:** Matthews Correlation Coefficient, **AUROC:** Area Under the Receiver Operating Characteristic curve, **SVC:** Support Vector Classifier, **ETree:** Extratree

**Table S22. The performance of different machine learning models developed using hybrid feature (Binary pattern+PSSM profile) with window length 5 on balanced dataset.**

| **Machine Learning Techniques**  (**Parameters**) | **Main Dataset** | | | | | **Validation Dataset** | | | | |
| --- | --- | --- | --- | --- | --- | --- | --- | --- | --- | --- |
|  | **Sen** | **Spc** | **Acc** | **MCC** | **AUROC** | **Sen** | **Spc** | **Acc** | **MCC** | **AUROC** |
| SVC  (g=, c=) | 77.72 | 75.98 | 76.85 | 0.54 | 0.84 | 70.98 | 75.91 | 73.45 | 0.47 | 0.79 |
| Random Forest  (Ntree=) | 77.89 | 77.83 | 77.86 | 0.56 | 0.86 | 73.58 | 77.46 | 75.52 | 0.51 | 0.83 |
| ExtraTree Classifier  (Ntree=) | 80.64 | 76.09 | 78.37 | 0.57 | 0.86 | 75.13 | 75.39 | 75.26 | 0.51 | 0.84 |
| K Nearest Neighbor  (n_neighbors=, algorithm=, weights=) | 73.79 | 72.45 | 73.12 | 0.46 | 0.82 | 68.39 | 74.35 | 71.37 | 0.43 | 0.80 |
| Multi Layer Perceptron  (activation=, solver=, hidden layer size=, maximum iteration=10) | 71.66 | 70.59 | 71.13 | 0.42 | 0.78 | 69.95 | 68.91 | 69.43 | 0.39 | 0.76 |
| Ridge Classifier  (alpha=1) | 73.40 | 70.48 | 71.94 | 0.44 | 0.79 | 74.35 | 63.99 | 69.17 | 0.39 | 0.76 |

*** Sen:** Sensitivity, **Spc:** Specificity, **Acc:** Accuracy, **MCC:** Matthews Correlation Coefficient, **AUROC:** Area Under the Receiver Operating Characteristic curve.

**Table S23. The performance of different machine learning models developed using hybrid feature (Binary pattern+PSSM profile) with window length 7 on balanced dataset.**

| **Machine Learning Techniques**  (**Parameters**) | **Main Dataset** | | | | | **Validation Dataset** | | | | |
| --- | --- | --- | --- | --- | --- | --- | --- | --- | --- | --- |
|  | **Sen** | **Spc** | **Acc** | **MCC** | **AUROC** | **Sen** | **Spc** | **Acc** | **MCC** | **AUROC** |
| SVC  (g=, c=) | 78.79 | 77.78 | 78.28 | 0.57 | 0.86 | 74.09 | 76.17 | 75.13 | 0.50 | 0.82 |
| Random Forest  (Ntree=) | 80.42 | 77.33 | 78.87 | 0.58 | 0.86 | 75.65 | 78.24 | 76.94 | 0.54 | 0.84 |
| ExtraTree Classifier  (Ntree=) | 79.91 | 78.62 | 79.26 | 0.59 | 0.87 | 73.83 | 79.79 | 76.81 | 0.54 | 0.85 |
| K Nearest Neighbor  (n_neighbors=, algorithm=, weights=) | 74.97 | 77.95 | 76.46 | 0.53 | 0.83 | 67.36 | 78.50 | 72.93 | 0.46 | 0.80 |
| Multi Layer Perceptron  (activation=, solver=, hidden layer size=, maximum iteration=10) | 75.36 | 72.11 | 73.74 | 0.47 | 0.81 | 62.44 | 80.83 | 71.63 | 0.44 | 0.78 |
| Ridge Classifier  (alpha=1) | 74.52 | 72.90 | 73.71 | 0.47 | 0.81 | 75.39 | 68.39 | 71.89 | 0.44 | 0.78 |

*** Sen:** Sensitivity, **Spc:** Specificity, **Acc:** Accuracy, **MCC:** Matthews Correlation Coefficient, **AUROC:** Area Under the Receiver Operating Characteristic curve.

**Table S24. The performance of different machine learning models developed using hybrid feature (Binary pattern+PSSM profile) with window length 9 on balanced dataset.**

| **Machine Learning Techniques**  (**Parameters**) | **Main Dataset** | | | | | **Validation Dataset** | | | | |
| --- | --- | --- | --- | --- | --- | --- | --- | --- | --- | --- |
|  | **Sen** | **Spc** | **Acc** | **MCC** | **AUROC** | **Sen** | **Spc** | **Acc** | **MCC** | **AUROC** |
| SVC  (g=, c=) | 80.70 | 78.06 | 79.38 | 0.59 | 0.87 | 76.17 | 75.91 | 76.04 | 0.52 | 0.84 |
| Random Forest  (Ntree=) | 79.07 | 77.95 | 78.51 | 0.57 | 0.87 | 74.87 | 77.98 | 76.42 | 0.53 | 0.84 |
| ExtraTree Classifier  (Ntree=) | 79.80 | 79.24 | 79.52 | 0.59 | 0.88 | 73.58 | 81.09 | 77.33 | 0.55 | 0.86 |
| K Nearest Neighbor  (n_neighbors=, algorithm=, weights=) | 76.71 | 80.13 | 78.42 | 0.57 | 0.85 | 69.95 | 78.50 | 74.22 | 0.49 | 0.81 |
| Multi Layer Perceptron  (activation=, solver=, hidden layer size=, maximum iteration=10) | 75.14 | 73.85 | 74.49 | 0.49 | 0.81 | 74.35 | 67.62 | 70.98 | 0.42 | 0.79 |
| Ridge Classifier  (alpha=1) | 74.86 | 73.12 | 73.99 | 0.48 | 0.82 | 74.87 | 67.88 | 71.37 | 0.43 | 0.79 |

*** Sen:** Sensitivity, **Spc:** Specificity, **Acc:** Accuracy, **MCC:** Matthews Correlation Coefficient, **AUROC:** Area Under the Receiver Operating Characteristic curve.

**Table S25. The performance of different machine learning models developed using hybrid feature (Binary pattern+PSSM profile) with window length 11 on balanced dataset.**

| **Machine Learning Techniques**  (**Parameters**) | **Main Dataset** | | | | | **Validation Dataset** | | | | |
| --- | --- | --- | --- | --- | --- | --- | --- | --- | --- | --- |
|  | **Sen** | **Spc** | **Acc** | **MCC** | **AUROC** | **Sen** | **Spc** | **Acc** | **MCC** | **AUROC** |
| SVC  (g=, c=) | 80.98 | 78.62 | 79.80 | 0.60 | 0.87 | 76.68 | 76.94 | 76.81 | 0.54 | 0.85 |
| Random Forest  (Ntree=) | 79.63 | 78.00 | 78.82 | 0.58 | 0.87 | 72.80 | 79.79 | 76.30 | 0.53 | 0.85 |
| ExtraTree Classifier  (Ntree=) | 82.49 | 76.37 | 79.43 | 0.59 | 0.88 | 77.72 | 80.05 | 78.89 | 0.58 | 0.87 |
| K Nearest Neighbor  (n_neighbors=, algorithm=, weights=) | 75.59 | 79.69 | 77.64 | 0.55 | 0.84 | 71.76 | 80.31 | 76.04 | 0.52 | 0.83 |
| Multi Layer Perceptron  (activation=, solver=, hidden layer size=, maximum iteration=10) | 75.08 | 74.41 | 74.75 | 0.49 | 0.82 | 73.58 | 74.87 | 74.22 | 0.48 | 0.81 |
| Ridge Classifier  (alpha=1) | 75.48 | 74.30 | 74.89 | 0.50 | 0.82 | 73.06 | 68.39 | 70.73 | 0.41 | 0.80 |

*** Sen:** Sensitivity, **Spc:** Specificity, **Acc:** Accuracy, **MCC:** Matthews Correlation Coefficient, **AUROC:** Area Under the Receiver Operating Characteristic curve.

**Table S26. The performance of different machine learning models developed using hybrid feature (Binary pattern+PSSM profile) with window length 13 on balanced dataset.**

| **Machine Learning Techniques**  (**Parameters**) | **Main Dataset** | | | | | **Validation Dataset** | | | | |
| --- | --- | --- | --- | --- | --- | --- | --- | --- | --- | --- |
|  | **Sen** | **Spc** | **Acc** | **MCC** | **AUROC** | **Sen** | **Spc** | **Acc** | **MCC** | **AUROC** |
| SVC  (g=, c=) | 81.82 | 78.00 | 79.91 | 0.60 | 0.88 | 77.20 | 79.02 | 78.11 | 056 | 0.86 |
| Random Forest  (Ntree=) | 79.97 | 78.23 | 79.10 | 0.58 | 0.87 | 75.39 | 80.31 | 77.85 | 0.56 | 0.85 |
| ExtraTree Classifier  (Ntree=) | 81.82 | 75.65 | 78.73 | 0.58 | 0.88 | 77.46 | 78.50 | 77.98 | 0.56 | 0.86 |
| K Nearest Neighbor  (n_neighbors=, algorithm=, weights=) | 75.65 | 80.86 | 78.25 | 0.57 | 0.85 | 68.39 | 80.05 | 74.22 | 0.49 | 0.81 |
| Multi Layer Perceptron  (activation=, solver=, hidden layer size=, maximum iteration=10) | 75.36 | 74.30 | 74.83 | 0.50 | 0.83 | 72.28 | 75.65 | 73.96 | 0.48 | 0.81 |
| Ridge Classifier  (alpha=1) | 75.36 | 75.08 | 75.22 | 0.50 | 0.82 | 72.80 | 70.73 | 71.76 | 0.44 | 0.80 |

*** Sen:** Sensitivity, **Spc:** Specificity, **Acc:** Accuracy, **MCC:** Matthews Correlation Coefficient, **AUROC:** Area Under the Receiver Operating Characteristic curve.

**Table S27. The performance of different machine learning models developed using hybrid feature (Binary pattern+PSSM profile) with window length 15 on balanced dataset.**

| **Machine Learning Techniques**  (**Parameters**) | **Main Dataset** | | | | | **Validation Dataset** | | | | |
| --- | --- | --- | --- | --- | --- | --- | --- | --- | --- | --- |
|  | **Sen** | **Spc** | **Acc** | **MCC** | **AUROC** | **Sen** | **Spc** | **Acc** | **MCC** | **AUROC** |
| SVC  (g=, c=) | 81.37 | 80.02 | 80.70 | 0.61 | 0.88 | 77.20 | 78.50 | 77.85 | 0.56 | 0.86 |
| Random Forest  (Ntree=) | 79.91 | 79.01 | 79.46 | 0.59 | 0.87 | 73.32 | 79.27 | 76.30 | 0.53 | 0.85 |
| ExtraTree Classifier  (Ntree=) | 79.69 | 80.58 | 80.13 | 0.60 | 0.88 | 73.32 | 82.12 | 77.72 | 0.56 | 0.87 |
| K Nearest Neighbor  (n_neighbors=, algorithm=, weights=) | 76.43 | 76.21 | 76.32 | 0.53 | 0.85 | 70.98 | 78.50 | 74.74 | 0.50 | 0.83 |
| Multi Layer Perceptron  (activation=, solver=, hidden layer size=, maximum iteration=10) | 75.53 | 73.68 | 74.61 | 0.49 | 0.83 | 70.47 | 76.42 | 73.45 | 0.47 | 0.82 |
| Ridge Classifier  (alpha=1) | 75.25 | 74.13 | 74.69 | 0.49 | 0.82 | 72.02 | 71.24 | 71.63 | 0.43 | 0.81 |

*** Sen:** Sensitivity, **Spc:** Specificity, **Acc:** Accuracy, **MCC:** Matthews Correlation Coefficient, **AUROC:** Area Under the Receiver Operating Characteristic curve.

**Table S28. The performance of different machine learning models developed using hybrid feature (Binary pattern+PSSM profile) with window length 17 on balanced dataset.**

| **Machine Learning Techniques**  (**Parameters**) | **Main Dataset** | | | | | **Validation Dataset** | | | | |
| --- | --- | --- | --- | --- | --- | --- | --- | --- | --- | --- |
|  | **Sen** | **Spc** | **Acc** | **MCC** | **AUROC** | **Sen** | **Spc** | **Acc** | **MCC** | **AUROC** |
| SVC  (g=, c=) | 81.54 | 79.91 | 80.72 | 0.61 | 0.88 | 78.76 | 77.72 | 78.24 | 0.56 | 0.87 |
| Random Forest  (Ntree=) | 80.13 | 79.12 | 79.63 | 0.59 | 0.87 | 74.61 | 78.50 | 76.55 | 0.53 | 0.86 |
| ExtraTree Classifier  (Ntree=) | 79.80 | 80.75 | 80.27 | 0.61 | 0.88 | 74.87 | 81.61 | 78.24 | 0.57 | 0.87 |
| K Nearest Neighbor  (n_neighbors=, algorithm=, weights=) | 82.21 | 68.69 | 75.45 | 0.51 | 0.85 | 79.79 | 72.80 | 76.30 | 0.53 | 0.84 |
| Multi Layer Perceptron  (activation=, solver=, hidden layer size=, maximum iteration=10) | 76.26 | 74.24 | 75.25 | 0.51 | 0.84 | 68.91 | 77.98 | 73.45 | 0.47 | 0.82 |
| Ridge Classifier  (alpha=1) | 75.25 | 75.08 | 75.17 | 0.50 | 0.82 | 70.98 | 72.02 | 71.50 | 0.43 | 0.81 |

*** Sen:** Sensitivity, **Spc:** Specificity, **Acc:** Accuracy, **MCC:** Matthews Correlation Coefficient, **AUROC:** Area Under the Receiver Operating Characteristic curve.

**Table S29. The performance of different machine learning models developed using hybrid feature (Binary pattern+PSSM profile) with window length 19 on balanced dataset.**

| **Machine Learning Techniques**  (**Parameters**) | **Main Dataset** | | | | | **Validation Dataset** | | | | |
| --- | --- | --- | --- | --- | --- | --- | --- | --- | --- | --- |
|  | **Sen** | **Spc** | **Acc** | **MCC** | **AUROC** | **Sen** | **Spc** | **Acc** | **MCC** | **AUROC** |
| SVC  (g=, c=) | 83.50 | 77.67 | 80.58 | 0.61 | 0.89 | 81.09 | 75.91 | 78.50 | 0.57 | 0.87 |
| Random Forest  (Ntree=) | 80.70 | 78.73 | 79.71 | 0.59 | 0.87 | 73.83 | 78.24 | 76.04 | 0.52 | 0.86 |
| ExtraTree Classifier  (Ntree=) | 80.08 | 81.09 | 80.58 | 0.61 | 0.88 | 74.35 | 80.31 | 77.33 | 0.55 | 0.86 |
| K Nearest Neighbor  (n_neighbors=, algorithm=, weights=) | 80.75 | 71.94 | 76.35 | 0.53 | 0.85 | 76.94 | 72.54 | 74.74 | 0.50 | 0.84 |
| Multi Layer Perceptron  (activation=, solver=, hidden layer size=, maximum iteration=10) | 78.11 | 75.20 | 76.66 | 0.53 | 0.84 | 76.17 | 73.06 | 74.61 | 0.49 | 0.82 |
| Ridge Classifier  (alpha=1) | 75.70 | 74.52 | 75.11 | 0.50 | 0.83 | 72.80 | 72.02 | 72.41 | 0.45 | 0.81 |

*** Sen:** Sensitivity, **Spc:** Specificity, **Acc:** Accuracy, **MCC:** Matthews Correlation Coefficient, **AUROC:** Area Under the Receiver Operating Characteristic curve.

**Table S30. The performance of different machine learning models developed using hybrid feature (Binary pattern+PSSM profile) with window length 21 on balanced dataset.**

| **Machine Learning Techniques**  (**Parameters**) | **Main Dataset** | | | | | **Validation Dataset** | | | | |
| --- | --- | --- | --- | --- | --- | --- | --- | --- | --- | --- |
|  | **Sen** | **Spc** | **Acc** | **MCC** | **AUROC** | **Sen** | **Spc** | **Acc** | **MCC** | **AUROC** |
| SVC  (g=, c=) | 83.00 | 76.94 | 79.97 | 0.60 | 0.88 | 79.53 | 76.42 | 77.98 | 0.56 | 0.87 |
| Random Forest  (Ntree=) | 80.53 | 77.72 | 79.12 | 0.58 | 0.87 | 75.39 | 77.98 | 76.68 | 0.53 | 0.86 |
| ExtraTree Classifier  (Ntree=) | 80.08 | 80.58 | 80.33 | 0.61 | 0.88 | 75.91 | 81.87 | 78.89 | 0.58 | 0.87 |
| K Nearest Neighbor  (n_neighbors=, algorithm=, weights=) | 79.46 | 72.56 | 76.01 | 0.52 | 0.86 | 73.06 | 75.65 | 74.35 | 0.49 | 0.83 |
| Multi Layer Perceptron  (activation=, solver=, hidden layer size=, maximum iteration=10) | 75.81 | 75.36 | 75.59 | 0.51 | 0.83 | 70.98 | 77.98 | 74.48 | 0.49 | 0.82 |
| Ridge Classifier  (alpha=1) | 75.98 | 73.79 | 74.89 | 0.50 | 0.82 | 72.02 | 75.13 | 73.58 | 0.47 | 0.81 |

*** Sen:** Sensitivity, **Spc:** Specificity, **Acc:** Accuracy, **MCC:** Matthews Correlation Coefficient, **AUROC:** Area Under the Receiver Operating Characteristic curve.

**Table S31. The performance of different machine learning models developed using hybrid feature (Binary pattern+PSSM profile) with window length 23 on balanced dataset.**

| **Machine Learning Techniques**  (**Parameters**) | **Main Dataset** | | | | | **Validation Dataset** | | | | |
| --- | --- | --- | --- | --- | --- | --- | --- | --- | --- | --- |
|  | **Sen** | **Spc** | **Acc** | **MCC** | **AUROC** | **Sen** | **Spc** | **Acc** | **MCC** | **AUROC** |
| SVC  (g=, c=) | 82.21 | 76.88 | 79.55 | 0.59 | 0.88 | 79.27 | 77.20 | 78.24 | 0.56 | 0.87 |
| Random Forest  (Ntree=) | 80.02 | 78.40 | 79.21 | 0.58 | 0.87 | 75.65 | 79.27 | 77.46 | 0.55 | 0.86 |
| ExtraTree Classifier  (Ntree=) | 80.86 | 79.35 | 80.11 | 0.60 | 0.88 | 74.87 | 82.38 | 78.63 | 0.57 | 0.87 |
| K Nearest Neighbor  (n_neighbors=, algorithm=, weights=) | 77.33 | 75.03 | 76.18 | 0.52 | 0.85 | 71.16 | 75.13 | 73.45 | 0.47 | 0.83 |
| Multi Layer Perceptron  (activation=, solver=, hidden layer size=, maximum iteration=10) | 76.26 | 74.69 | 75.48 | 0.51 | 0.83 | 65.28 | 83.16 | 74.22 | 0.49 | 0.82 |
| Ridge Classifier  (alpha=1) | 75.31 | 73.57 | 74.44 | 0.49 | 0.82 | 73.06 | 75.39 | 74.22 | 0.48 | 0.81 |

*** Sen:** Sensitivity, **Spc:** Specificity, **Acc:** Accuracy, **MCC:** Matthews Correlation Coefficient, **AUROC:** Area Under the Receiver Operating Characteristic curve.

**Table S32. The performance of PSSM profile based models developed using different machine learning techniques on additional dataset at window size 17.**

| **Machine Learning Techniques**  (**Parameters**) | **Additional Dataset** | | | | |
| --- | --- | --- | --- | --- | --- |
|  | **Sen** | **Spc** | **Acc** | **MCC** | **AUROC** |
| SVC_MaxMCC#_  (g=0.01, c=5) | 41.48 | 99.89 | 97.14 | 0.62 | 0.88 |
| SVC_Balanced*_  (g=0.01, c=5) | 81.48 | 82.31 | 82.27 | 0.33 | 0.88 |
| Random Forest _MaxMCC#_  (Ntree = 700) | 42.96 | 99.30 | 96.65 | 0.55 | 0.87 |
| Random Forest _Balanced*_  (Ntree = 700) | 77.78 | 76.01 | 76.10 | 0.26 | 0.87 |
| Extra Tree _MaxMCC#_  (Ntree = 1000) | 45.19 | 99.38 | 96.82 | 0.58 | 0.86 |
| Extra Tree _Balanced*_  (Ntree = 1000) | 77.04 | 74.62 | 74.74 | 0.24 | 0.86 |
| K Nearest Neighbors _MaxMCC#_  (neighbors=10, algorithm=auto, weights=distance) | 54.81 | 98.65 | 96.58 | 0.59 | 0.83 |
| K Nearest Neighbors _Balanced*_  (neighbors=10, algorithm=auto, weights=distance) | 74.07 | 81.84 | 81.47 | 0.29 | 0.83 |
| Muti Layer Perceptron _MaxMCC#_  (activation=relu, solver=adam, hidden_layer_sizes=21, max_iter=10) | 7.41 | 100.00 | 95.64 | 0.27 | 0.82 |
| Muti Layer Perceptron _Balanced*_  (activation=relu, solver=adam, hidden_layer_sizes=21, max_iter=10) | 75.65 | 71.15 | 71.35 | 0.21 | 0.82 |
| Ridge Classifier _MaxMCC#_  (alpha=1) | 8.89 | 99.85 | 95.57 | 0.25 | 0.78 |
| Ridge Classifier _Balanced*_  (alpha=1) | 71.85 | 69.50 | 69.61 | 0.19 | 0.78 |

**Note: Sen:** Sensitivity, **Spc:** Specificity, **Acc:** Accuracy, **MCC:** Matthews Correlation Coefficient, **AUROC:** Area Under the Receiver Operating Characteristic curve.

***:** Results are reported on balanced sensitivity and specificity; **#:** Results are reported for which maximum MCC was obtained.

**Figure S1.** Normalized propensity score of SAM interacting and non-interacting residues.

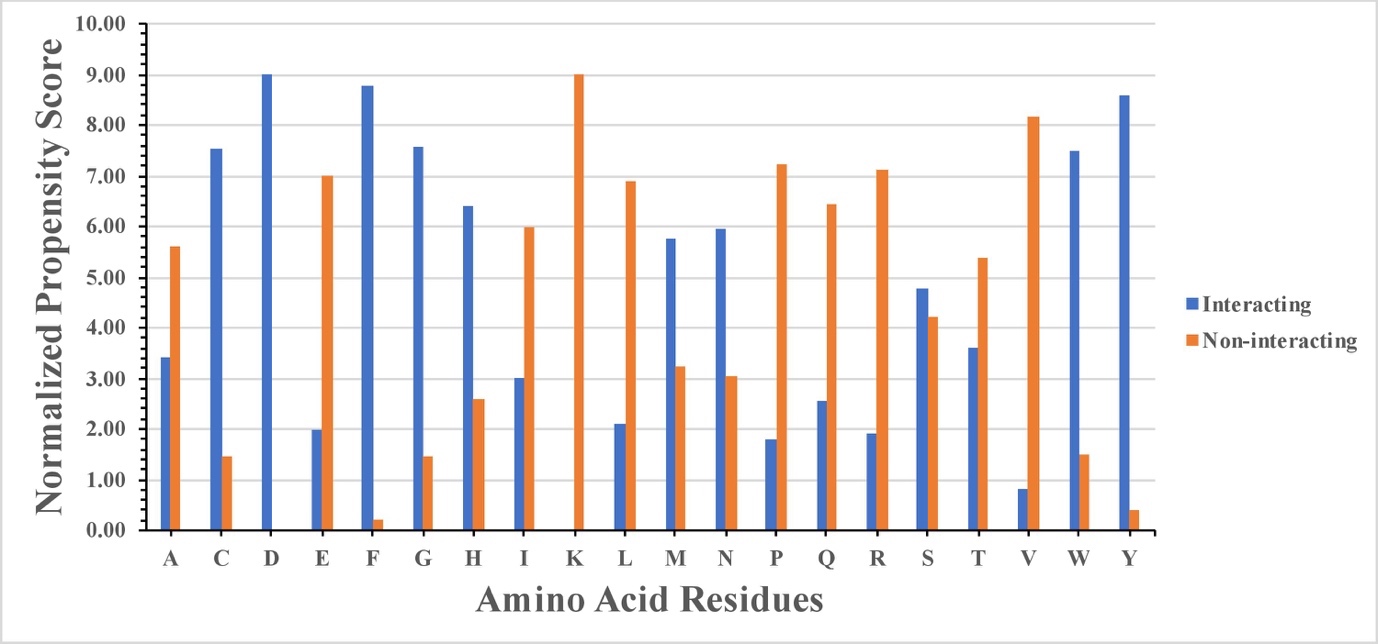

**Figure S2.** Percent composition of various physiochemical properties of SAM interacting and non-interacting residues.

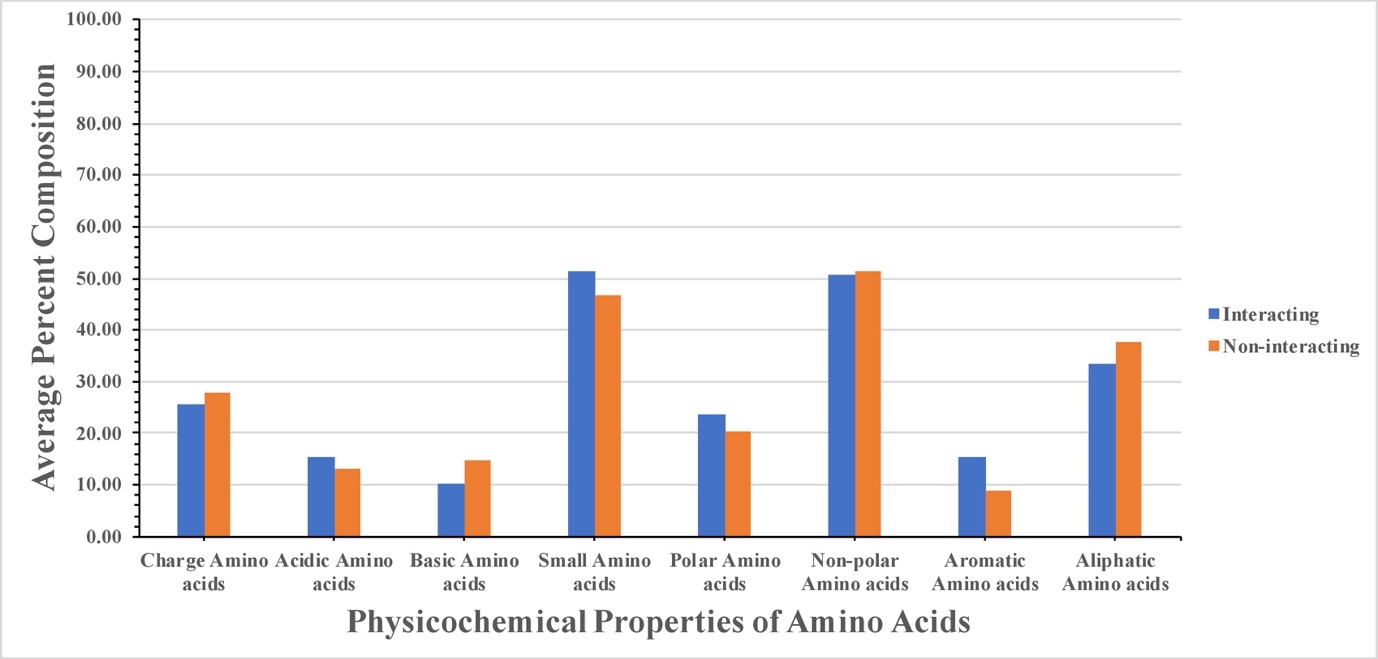

**Figure S3.** AUROC plots obtained for various window length developed using hybrid feature on balanced dataset for (a) training dataset and (b) validation dataset.

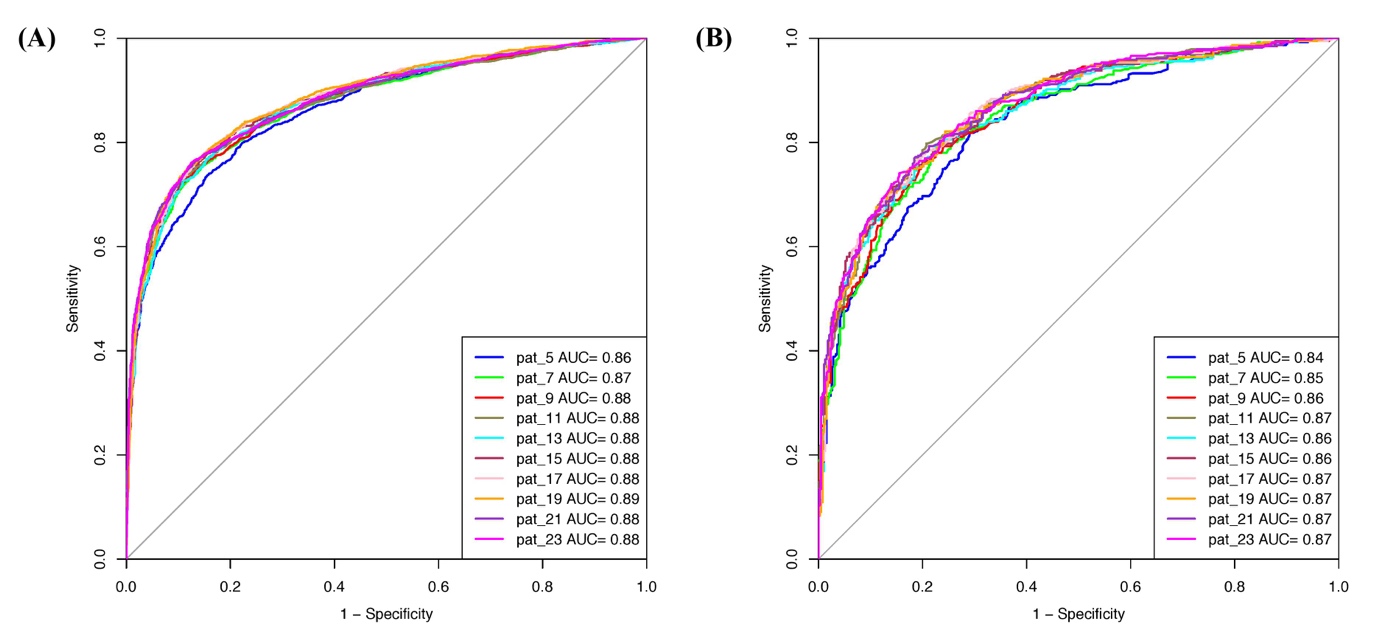
